# A pentameric protein ring with novel architecture is required for herpesviral packaging

**DOI:** 10.1101/2020.07.16.206755

**Authors:** Allison L. Didychuk, Stephanie N. Gates, Matthew R. Gardner, Lisa M. Strong, Andreas Martin, Britt A. Glaunsinger

**Affiliations:** Department of Plant and Microbial Biology, University of California, Berkeley, California, USA; Department of Molecular and Cell Biology, University of California, Berkeley, California, USA; California Institute for Quantitative Biosciences, University of California, Berkeley, California 94720, USA; Howard Hughes Medical Institute, University of California, Berkeley, California, USA

**Author notes:** Correspondence should be addressed to Britt Glaunsinger.

## Abstract

Genome packaging in large double-stranded DNA viruses requires a powerful molecular motor to force the viral genome into nascent capsids. This process appears mechanistically similar in two evolutionarily distant viruses, the herpesviruses and the tailed bacteriophages, which infect different kingdoms of life. While the motor and mechanism as a whole are thought to be conserved, accessory factors that influence packaging are divergent and poorly understood, despite their essential roles. An accessory factor required for herpesviral packaging is encoded by ORF68 in the oncogenic virus Kaposi’s sarcoma-associated herpesvirus (KSHV), whose homolog in Epstein Barr Virus (EBV) is BFLF1. Here, we present structures of both KSHV ORF68 and EBV BFLF1, revealing that these proteins form a highly similar homopentameric ring. The central channel of this ring is positively charged, and we demonstrate that this region of KSHV ORF68 binds double-stranded DNA. Mutation of individual positively charged residues within but not outside the channel ablates DNA binding, and in the context of KSHV infection these mutants fail to package the viral genome or produce progeny virions. Thus, we propose a model in which ORF68 facilitates the transfer of newly replicated viral genomes to the packaging motor.

---

Herpesviruses are large double-stranded DNA viruses that cause a variety of diseases in humans. The ability of herpesviruses to efficiently evade the immune system and establish latency, coupled with few available treatments and vaccines, means that nearly all adults in the world harbor at least one of the nine human herpesviruses. Herpes simplex virus type 1 (HSV-1) is an alphaherpesvirus that causes cold sores and genital sores. Human cytomegalovirus (HCMV) is a betaherpesvirus that can cause mononucleosis and congenital birth defects. The human gammaherpesviruses Kaposi’s sarcoma-associated virus (KSHV) and Epstein-Barr virus (EBV) are oncogenic viruses, causing cancers such as primary effusion lymphoma (PEL) and Kaposi’s sarcoma (in the case of KSHV).

Despite 400 million years of evolution separating the human herpesviruses, several core pathways in replication are conserved^1^. Near the end of the lytic cycle, herpesviruses replicate their genome as a head-to-tail concatemer of linked genomes separated by terminal repeats. Cleavage to produce a unit-length genome is intimately tied to packaging and occurs only after that genome is successfully transferred into a capsid. DNA packaging in the tailed bacteriophages is thought to be mechanistically similar to that of herpesviruses^2^. Despite infecting hosts in different kingdoms, both groups of viruses use an icosahedral capsid and an architecturally similar portal protein through which DNA is packaged^2,3^. Furthermore, both depend on a “terminase” motor responsible for packaging and cleavage of the genome. The large subunit of the terminase is the most conserved gene across the herpesviruses and possesses sequence and structural similarity to phage terminases, supporting the hypothesis that packaging occurs through an evolutionarily ancient mechanism^2,4,5^. Packaging minimally requires recognition of the viral genome by the terminase, docking of the terminase-bound genome at the portal of a nascent capsid, translocation of the genome into the capsid by the terminase, and cleavage to release the remaining unpackaged concatemeric genome.

Cleavage and packaging in the herpesviruses, best studied in HSV-1, requires six conserved proteins in addition to the nascent capsid and concatemeric genome^6^: HSV-1 UL6, UL15, UL17, UL28, UL32, and UL33. Three of these proteins (UL15/UL28/UL33) form the terminase motor, and the portal protein is composed of a dodecamer of UL6^7,8^. UL17 encodes a capsid vertex-specific protein important for stabilizing the capsid^9–11^. In contrast, despite observations that UL32 and its homologs in HCMV (UL52) and KSHV (ORF68) are essential for production of packaged virions, its function in packaging remains unknown^12–15^. Phages lack an identifiable homolog of UL32 or ORF68, suggesting that an additional level of complexity exists in herpesvirus packaging.

Here, we applied a combination of structural biology and biochemistry to better define the role of the essential accessory protein ORF68 in KSHV packaging. We reveal the structure of KSHV ORF68 and its homolog in EBV (BFLF1), which adopt a novel fold and assemble into a homopentameric ring. The similarity of these structures, combined with homology modeling and negative stain electron microscopy of homologs from HSV-1 and HCMV, suggest that this topology is conserved across the *Herpesviridae.* The central channel of ORF68 is lined by positively charged residues that are necessary for nucleic acid binding and production of infectious virions. We hypothesize that the viral genome is threaded through the ORF68 ring, and that ORF68 acts as a scaffold on which the terminase assembles for genome packaging.

## RESULTS

### ORF68 forms a homopentameric ring

To structurally characterize KSHV ORF68, we purified the full-length protein from transiently transfected HEK293T cells. In agreement with our prior observation that ORF68 forms a multimer *in vitro*^13^, negative stain EM revealed rings comprised of five subunits (**Supplementary Fig. S1a**). A cryo-EM reconstruction of the pentamer was determined to 3.37 Å, from which an alanine backbone model was built (**Supplementary Fig. S1b**). However, as ORF68 bears no sequence homology to proteins outside of the *Herpesviridae, de novo* modeling was challenging. We subsequently crystallized ORF68 and solved its structure by molecular replacement with the initial cryo-EM model. Representative diffraction data are presented in **Supplementary Fig. S2,** and data collection and structure refinement statistics are listed in **Supplementary Table 1** (cryo-EM) and **Supplementary Table 2** (X-ray). The majority of the protein could be built in the X-ray structure, except for two disordered loops (residues 64-67 and residues 241-262), and a region in the central channel (residues 169-172 and 179-188). These regions were similarly disordered in the cryo-EM maps, suggesting inherent flexibility.

ORF68 forms a homopentameric ring with a diameter of ~120 Å (**Fig. 1a**). Each monomer contains three zinc fingers and several bundles of α-helices connected by loops. A DALI search^16^ identified proteins that are similarly α-helical in nature, but none with globally similar structures, suggesting that ORF68 adopts a novel fold. The conserved residues in ORF68 are generally located in hydrophobic regions, suggesting that they play structural roles (**Supplementary Fig. S3**). On the “top” face of the ring lies a short semi-structured loop (residues P27-N36), which represents a region where large insertions are observed in alpha- and beta-herpesviruses homologs (**Supplementary Fig. S4**). The central channel is constricted toward the top of the ring, with a width of ~25 Å, and widens to ~45 Å at the bottom (**Fig. 1a**). Two segments of each ORF68 monomer directly face this central channel: residues 167-188 and 435-451. Residues 435-451 form an α-helix that is anchored by H452, which coordinates a Zn ion at the bottom of the ring (**Fig. 1b**). Interestingly, residues 167-188 are largely disordered, despite being anchored by C191 and C192 that participate in coordination of the same Zn ion.

**Fig. 1.**
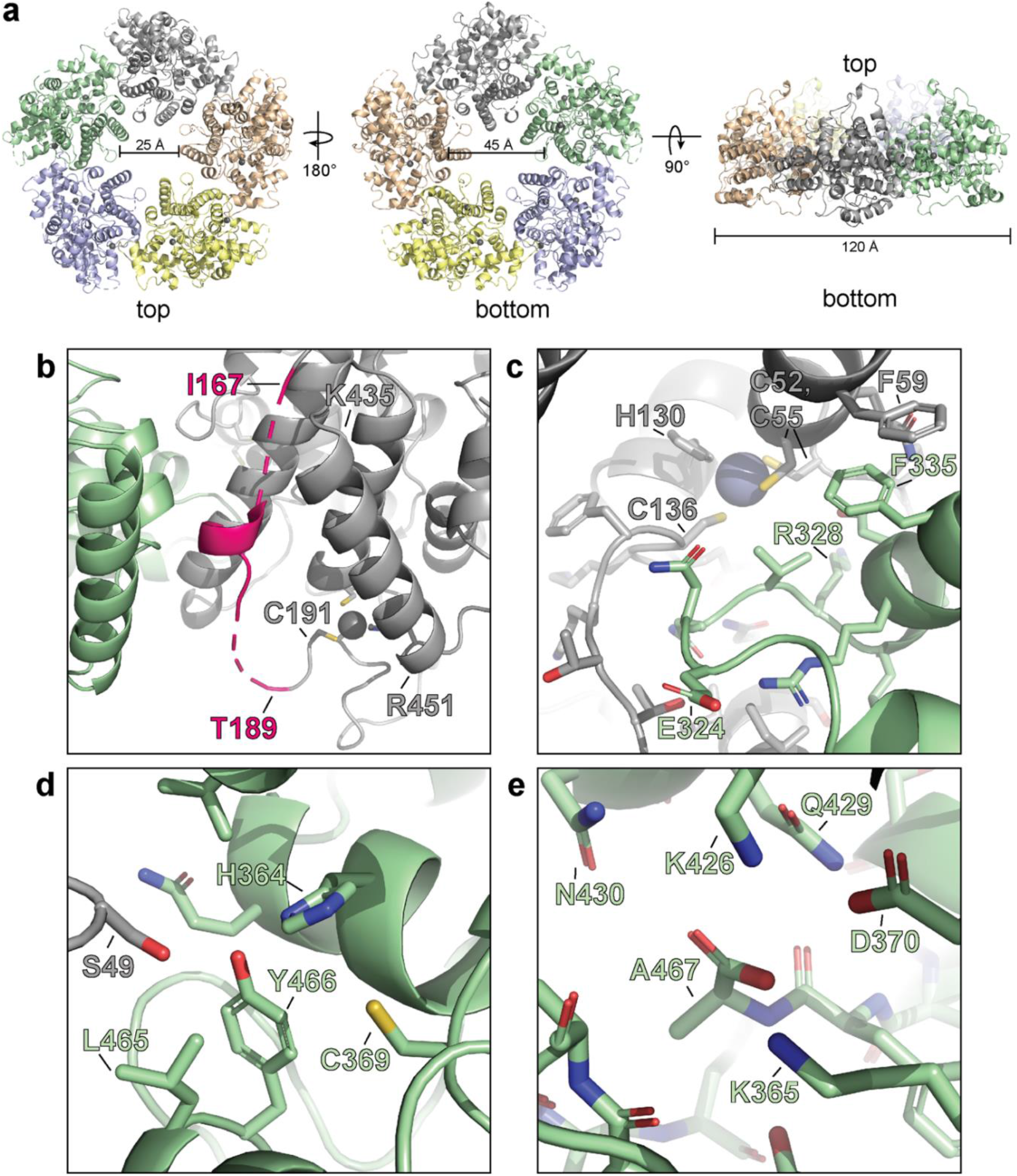
ORF68 forms a homopentameric ring. **a**, View from the top, bottom, and side of the ORF68 crystal structure. Pore size and overall diameter are highlighted. **b**, Residues 167-188 and 435-451 span the pore. Residues 435-451 form an α-helix, whereas residues 177-188 are largely disordered (highlighted in pink). Both regions are anchored by a zinc finger consisting of C191/C192/C415/H452. **c**, Subunit interface within the ORF68 pentamer, with one monomer shown in green and its clockwise neighbor in grey. **d**, ORF68’s penultimate residue, Y466, is buried near conserved residues C369 and H364, along with S49 from a neighboring monomer. **e**, The C-terminus of the protein (A467) is surrounded by conserved residues, including K365, D370, K426, and Q429.

ORF68 is stable as a pentamer even at high ionic strength (1 M NaCl, data not shown). The subunit interface includes ~1200 Å2 of buried surface area, consisting largely of van der Waals contacts and a stacking interaction between F59 of one subunit and F335 of the adjacent subunit. This interface is generally poorly conserved, with the exception of N150 and D331, which interact with the backbone of neighboring subunits. A loop consisting of residues 324-329 inserts into an adjacent monomer to interact with residues 136-145 and 55-59, an interaction that appears to be stabilized by the first zinc finger (**Fig. 1 c**).

The C-terminal tail of ORF68 is buried near the subunit interface. The penultimate residue, Y466, is perfectly conserved in all homologs and is buried near the highly conserved residues H364 and C369 (**Fig. 1d, Supplementary Fig. S4**). Furthermore, the C-terminal carboxyl group is surrounded by the highly conserved residues K365, D370, K426, and Q429 (**Fig. 1e**). We found that addition of a C-terminal tag reduces expression of the protein and prevents infectious virion production (data not shown). This effect was also observed in HCMV, where C-terminal tagging of the ORF68 homolog, UL52, prevented virion production and led to an aberrantly disperse localization throughout the cell^15^. Thus, disrupting the coordination of ORF68’s C-terminus at the subunit interface through addition of a tag likely interferes with pentamer formation, destabilizes the protein, and prevents its function in DNA packaging and cleavage.

### Zinc fingers in ORF68 and its homologs are necessary for stability

ORF68 contains several motifs with highly conserved cysteines and histidines. These residues form three CCCH-type zinc fingers, as supported by the tetrahedral coordination geometry, a large anomalous scattering signal, and C/H composition of the putative zinc fingers, along with the previous observation that the homolog from HSV-1, UL32, binds zinc^17^. The residues that comprise two of these zinc fingers (residues C52, C55, H130, and C136 in the first zinc finger; residues C296, C299, H366, and C373 in the second zinc finger) are perfectly conserved in all identified homologs of ORF68 (**Fig. 2a, b**, **Supplementary Fig. S4**). The third zinc finger (consisting of C191, C192, C415, and H452), is generally conserved in the alpha- and gammaherpesviruses (with the exception of the closely related model murine gammaherpesvirus MHV68), but is missing in the betaherpesviruses (**Fig. 2c**, **Supplementary Fig. S4**). This third zinc finger is also atypical in that two coordinating residues, C191 and C192, are adjacent to each other. Such noncanonical cysteine organization in a zinc finger is rare, but has also been observed in the structure of RPB10, a subunit of RNA polymerase^18^. There is additional weak anomalous density for a potential fourth metal binding site, coordinated by C169, C172, and H464, yet this was not included in the final model due to inconsistent density and stereochemistry.

**Fig. 2.**
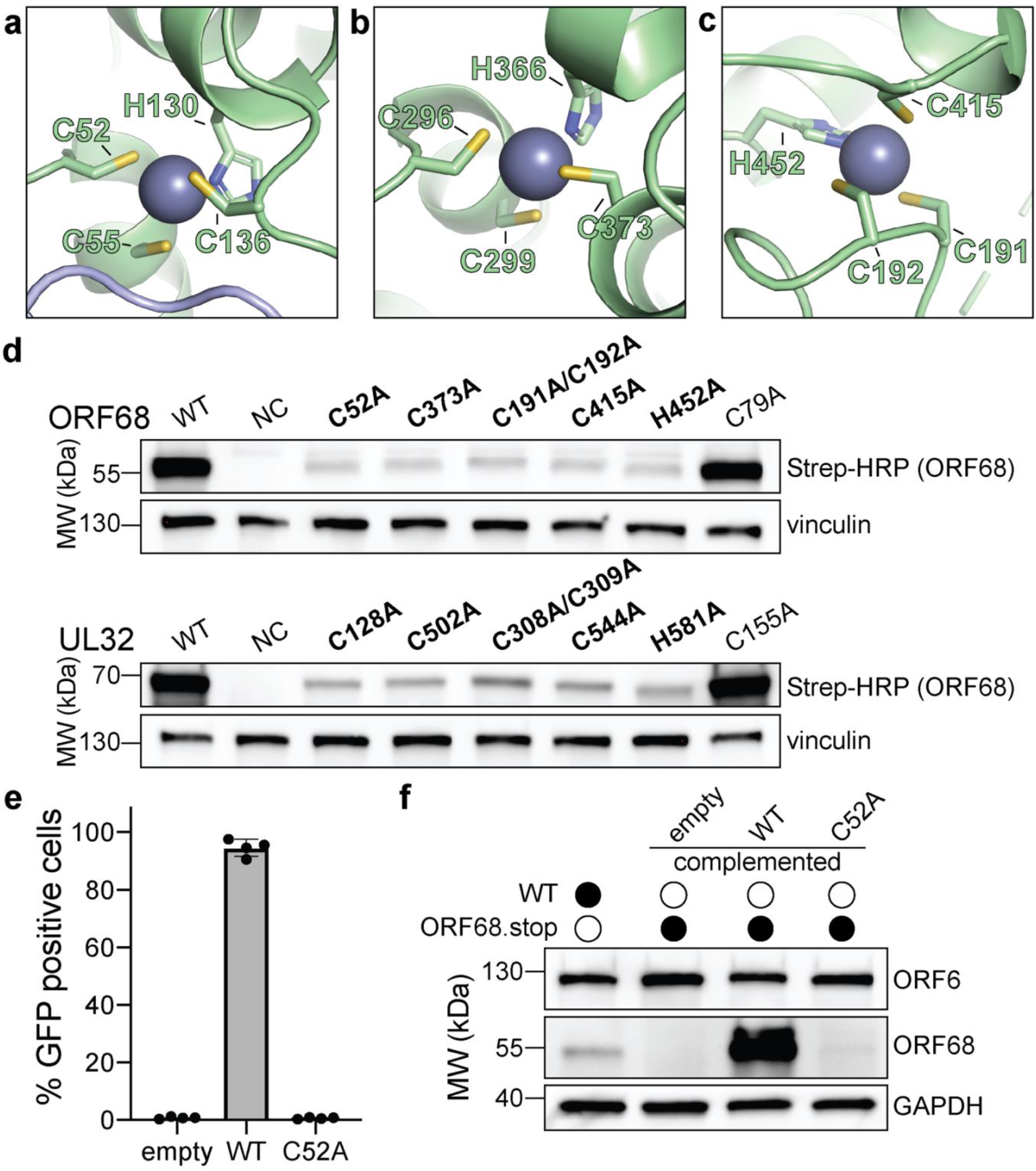
ORF68 and homologs are zinc-finger containing proteins. **a-c**, The Zn_2+_ ion within the three zinc finger motifs is shown as a blue sphere, while coordinating cysteines and histidines are shown in sticks. **d**, Western blot of whole cell lysate (33 μg) from HEK293T cells that were transfected with plasmids encoding wild-type and mutant variants of ORF68 (top) or UL32 (bottom), where residues involved in zinc finger coordination were mutated to alanine. Vinculin serves as a loading control. **e**, ORF68.stop iSLK cells were lentivirally transcomplemented with empty vector or with plasmids encoding wild-type or C52A ORF68. Progeny virion production by these cell lines was assayed by supernatant transfer and flow cytometry of target cells. **f**, Western blot of transcomplemented ORF68.stop iSLK cells used in **e**.

To evaluate the role of residues in Zn coordination, we generated ORF68 mutants containing cysteine to alanine substitutions within the zinc fingers or for a similarly conserved cysteine outside of the zinc finger motifs (residue C79) (**Fig. 2d**). ORF68 variants with a mutation in any of the zinc fingers were poorly expressed in transfected HEK293T cells, suggesting that the zinc fingers are required for structural stability of the protein. Several conserved cysteines in UL32 were previously identified as important for virion production in HSV-1^12^. We observed that these cysteines in UL32 are homologous to the zinc finger residues in ORF68 and are similarly essential for structural stability, as their substitution with alanine resulted in lower UL32 protein levels, whereas mutation of a conserved cysteine outside of the zinc fingers had no effect (**Fig. 2d**). Thus, UL32 likely also contains zinc fingers that are required for its structural stability.

We next tested whether these stabilizing zinc fingers in ORF68 were required for the production of infectious virions. Using a latently infected inducible SLK (iSLK) BAC16 cell line in which the KSHV genome contains two premature stop codons to prevent ORF68 expressions^13^, we assessed whether complementation with constitutively expressed ORF68 WT or the C52A zinc-finger mutant allowed for production of infectious virions in a supernatant transfer assay. Wild-type ORF68-expressing cells were able to produce virions sufficient to infect nearly 100% of target cells, while ORF68-C52A expressing cells were unable to produce progeny virions (**Fig. 2e**). ORF68-C52A could not be detected by western blot, suggesting that the C52A mutation is severely disruptive to the structure of ORF68 even in the context of viral infection (**Fig. 2f**).

### Homologs of ORF68 form similar structures

ORF68 homologs can be found in all known members of the *Herpesviridae*, although BLAST cannot identify candidates in the *Alloherpesviridae* and *Malacoherpesviridae,* related families in the order *Herpesvirales* that infect fish, amphibians, and mollusks. Homologs of ORF68 (467 residues) in the *Herpesviridae* range in size from 437 residues (MHV68 mu68) to 668 residues (HCMV UL52) and show generally low sequence conservation. Most of the differences in length come from N-terminal extensions or insertions in regions expected to be surface-exposed loops, suggestive of a conserved core structure (**Supplementary Fig. S4**).

We sought to determine whether ORF68 homologs have a conserved structure, which may help identify features important for their function in packaging. We purified BFLF1, the ORF68 homolog in EBV, from transiently transfected HEK293T cells. Interestingly, negative stain EM revealed that BFLF1 forms decameric rings, formed by two stacked pentamer rings (**Supplementary Fig. S1c**). We determined the structure of BFLF1 by cryo-EM at 3.60 Å resolution and found that it forms pentameric rings comparable in size to those of ORF68 and with a highly similar structure (**Fig. 3a, Supplementary Fig. S1d**). Based on the behavior in size-exclusion chromatography, the pentameric ring likely represents the active form for BFLF1. Its structure is highly similar to that of ORF68, with an rmsd of 1.09 between monomers and an overall rmsd of 1.95 for the complex (**Fig. 3a, b**). The internal core of the protein, including the first two zinc fingers (residues C54/C57/H132/C138 and C316/C319/H386/C393 in BFLF1), is highly conserved. Small variations occur in surface-exposed loops. Residues 180-240 (corresponding to residues 178-219 in ORF68) and 264-278 (corresponding to residues 244-258 in ORF68) could not be resolved, suggesting inherent flexibility. This flexibility also prevented unambiguous modeling of the putative third zinc finger, although BFLF1 residues C432 and H469 can be modeled, and C214/C215 likely lie within the immediately surrounding disordered region.

**Fig. 3.**
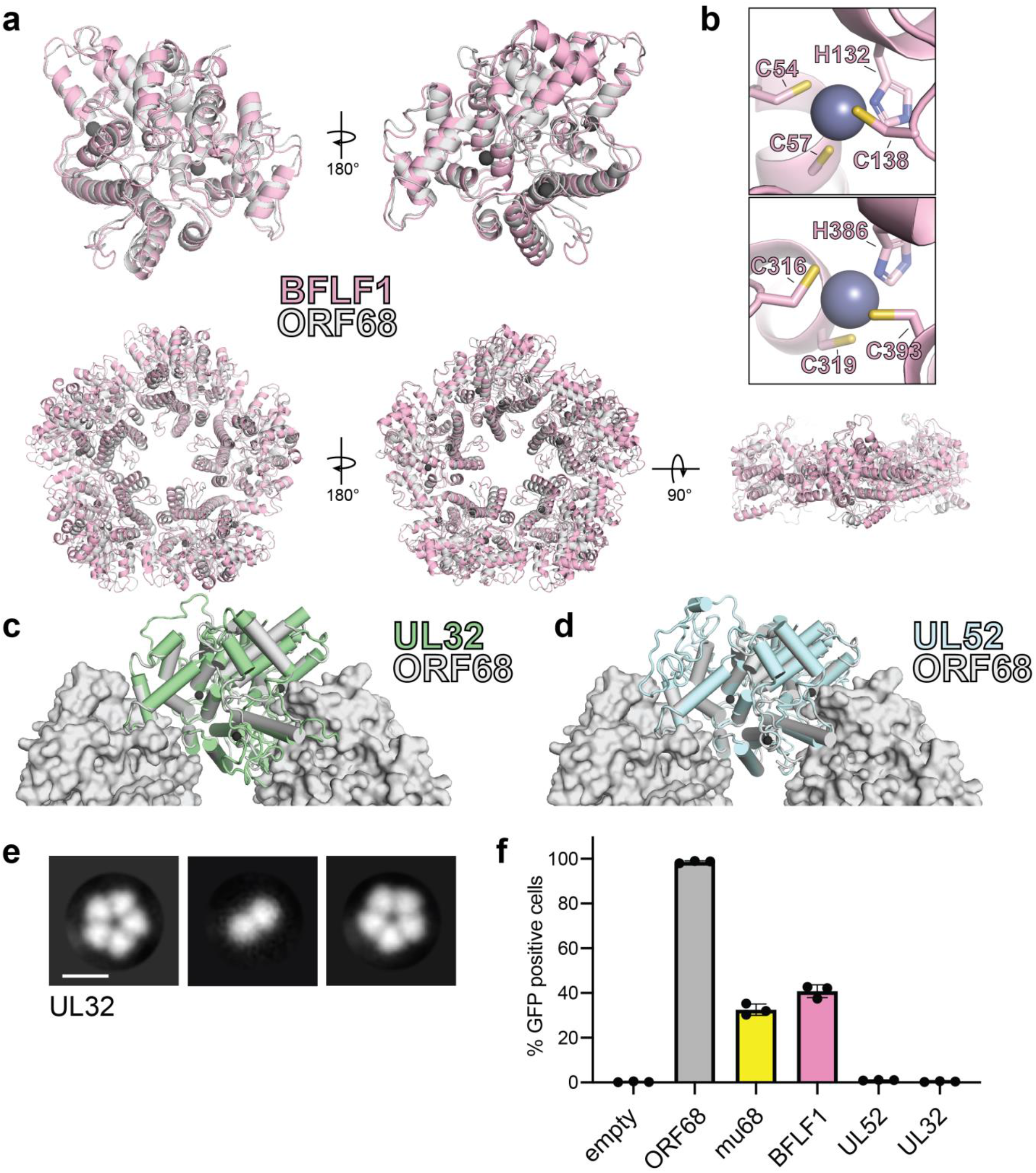
Homologs of ORF68 possess similar structures. **a**, Overlay of ORF68 (grey) and BFLF1 (pink) monomers (top) and their homopentameric complexes (bottom). **b**, BFLF1 contains at least two zinc fingers, with Zn2+ shown in grey. **c-d**, Homology modelling of UL32 (green) and UL52 (blue) suggest they have a highly similar structure to ORF68 (grey), with differences in surface-exposed loops and in the monomer-monomer interface. **e**, Representative 2D class averages from negative-stain EM of UL32. Scale bar = 100 Å. **f**, ORF68.stop iSLK cells were lentivirally transcomplemented with plasmids encoding N-terminally Strep-tagged ORF68 or homologs from EBV (BFLF1), MHV68 (mu68), HCMV (UL52), or untagged HSV-1 (UL32). Progeny virion production by these cell lines was assayed by supernatant transfer and flow cytometry of target cells.

Based on the high structural similarity between ORF68 and BFLF1, as well as the conservation of their core residues (**Supplementary Fig. S4**), we used the ORF68 structure with the SWISS-MODEL server^19^ to generate homology models for UL32 (from HSV-1) and UL52 (from HCMV) (**Fig. 3c, d**). These models suggest that the internal core of the protein is highly structurally conserved, and that the primary differences arise from surface-exposed loops that vary in length (**Supplementary Fig. S4**). We performed negative stain EM on UL32 to determine if it also retains homopentameric quaternary structure (**Fig. 3e**). Despite the low sequence homology to residues identified at the subunit interface of ORF68 and BFLF1, ring-shaped structures were observed for UL32 as well, and class averages suggested that it forms a homopentamer. Pentameric oligomerization is therefore a common feature of ORF68 and its homologs.

Next we investigated if homologs of ORF68 were sufficiently similar to functionally replace its essential role in viral packaging in KSHV. We generated stable cell line derivatives that constitutively expressed either ORF68 or its homolog from EBV (BFLF1), MHV68 (mu68), HCMV (UL52), or HSV-1 (UL32) to complement the stop-codon containing ORF68 in the KSHV genome (**Fig. 3f**). As expected, ORF68.stop cells complemented with ORF68 allowed for production of virions that infected nearly 100% of target cells, while complementation with empty vector failed to produce infectious virions. Homologs from the gammaherpesviruses, BFLF1 and mu68, were able to partially complement loss of ORF68, although not to levels comparable to ORF68-expressing cells. In contrast, homologs from the more distantly related alpha- and betaherpesviruses, UL32 and UL52, failed to complement deletion of ORF68. BFLF1 and mu68 are most similar to ORF68 (35% identity/52% similarity and 33% identity/50% similarity, respectively), while UL32 and UL52 are more diverged in sequence (25% identity/34% similarity and 22% identity/34% similarity respectively). Thus, although homologs across the herpesviruses share a common core fold and homopentameric architecture, other features or the identity of surface residues likely play a role for their function and the interaction with other components during DNA packaging.

### ORF68 binds dsDNA via its positively charged central channel

The calculated electrostatic surface of ORF68 shows a striking colocalization of positive charges on one side of the ring around the entrance of the central channel (**Fig. 4a**). We previously observed that ORF68 can bind an 800 bp dsDNA probe corresponding to the GC-rich terminal repeat of KSHV13, and therefore hypothesized that the electrostatic surface around the central channel of ORF68 could be important for dsDNA binding. Although we also previously observed that ORF68 has weak nuclease activity, we were unable to identify a motif suggestive of such activity in the structure. ORF68 binds 10-20 bp dsDNA with high affinity, while multiple binding events are observed on longer probes (**Supplementary Fig. S5a**, ref. 13). We selected a series of surface-exposed positively charged residues (arginine or lysine) on either side and throughout the central channel to test if substitution to alanine reduced dsDNA binding. All ORF68 mutants expressed and purified similar to wild-type ORF68 (**Supplementary Fig. S6a**), and were characterized in electrophoretic mobility shift assays regarding their dsDNA-binding affinity relative to wild-type ORF68 (**Fig. 4b, c**, **Supplementary Fig. S6b**). Mutations K435A (top, more constricted side of the central channel), R443A (middle of the central channel), and K174A/R179A/K182A (henceforth referred to as “3+”, in the disordered region of the central channel) all resulted in a drastic reduction in binding affinity, with negligible interactions even at 4 μM ORF68. The K450A/R451A mutation (bottom of the central channel) bound with lower affinity than wild-type ORF68, as indicated by the lack of concrete bands and a consequential smear. In contrast, mutation in two sets of surface-exposed positive residues located outside the channel, R14A/K310A and K395A/K396A, bound dsDNA comparably to wild-type ORF68 with a *Kd* of ~25 nM. Thus, while charge mutations on the periphery of the ring have no effect, mutations within the channel – and specifically near the more constricted top portion – are deleterious to dsDNA binding, likely because residues in and near the central channel from all subunits form a large binding interface with high charge density.

**Fig. 4.**
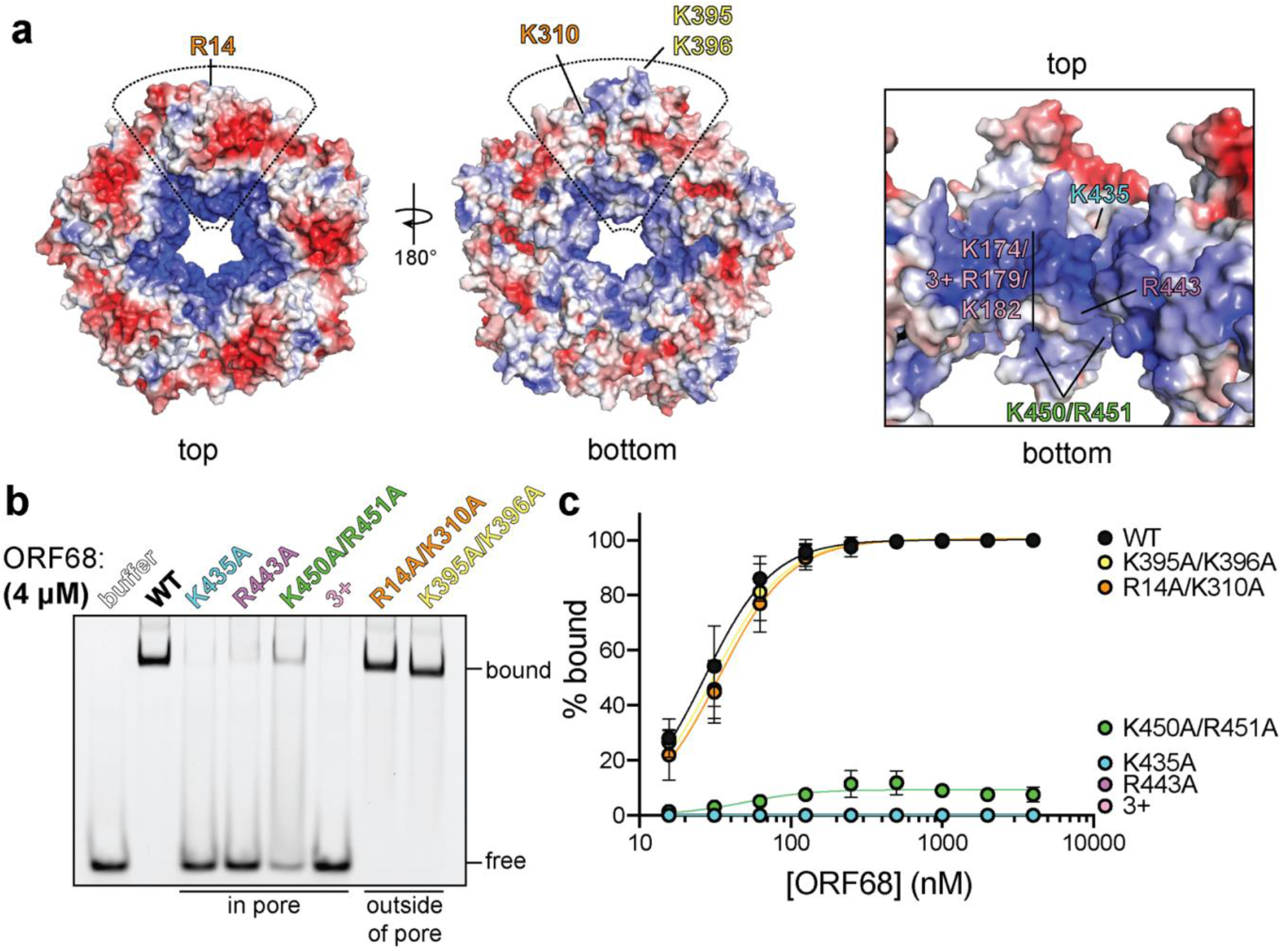
ORF68 binds nucleic acid in vitro via its central channel. **a**, Electrostatic surface of the ORF68 pentamer, contoured from +5 kT/e (blue) to −5 kT/e (red) and shown from the top (left), bottom (middle), and through the pore (right). The electrostatic surface lacks regions that were disordered in the structure, including residues 169-172 and 179-188, which face the central channel. The location of residues selected for mutation are indicated on one monomer of the pentamer. **b**, Electrophoretic mobility shift assay using fluorescein-labeled 20 bp dsDNA probe (10 nM) and wild-type or mutant ORF68 (4 μM). **c**, Binding curves for wild-type and mutant ORF68 interacting with the 20 bp dsDNA probe were determined by electrophoretic mobility shift assays as in **b**. Data represent the mean ± s.d. of three independent experiments. Data were fit with a nonlinear regression o the Hill equation.

Herpesviral DNA packaging requires both sequence specific and nonspecific DNA binding. Site-specific binding and cleavage within the terminal repeats is required to ensure full-length genomes are packaged; however, this role is thought to be fulfilled by the terminase^20,21^. Packaging also depends on non-sequence specific interactions with various factors, such as HSV-1 UL25 (KSHV ORF19) that binds DNA and has recently been shown to represent the “portal cap” that prevents DNA escape after packaging has been completed^9,11,22^. To assess the sequence specificity of ORF68, we compared its binding to a 20 bp sequence derived from the terminal repeats with high (85%) GC content versus a scrambled sequence with 50% GC content (**Supplementary Fig. 5b**). ORF68 bound with ~25 nM affinity to both substrates, suggesting that it recognizes dsDNA *in vitro* in a non-sequence-specific manner.

### Positively charged residues within the central channel are required for cleavage and packaging

We next sought to determine whether electrostatically-mediated nucleic acid binding is important during viral replication. Using the Red recombinase system^23^, we incorporated two ORF68 variants that ablate dsDNA binding *in vitro,* K435A and the 3+ mutant, as well as the corresponding mutant rescue (MR) control constructs with revertant mutations into the KSHV BAC16 genome (**Supplementary Fig. S7a, b**). We established latently infected iSLK cell lines harboring KSHV with a mutant copy of ORF68 and assessed their ability to produce infectious virions using the supernatant transfer assay (**Fig. 5a**). As expected, wild-type KSHV was able to infect nearly 100% of target cells, whereas the ORF68.stop cells failed to produce infectious virions. The ORF68-K435A cells had a pronounced virion-production defect, leading to infection of <5% of target cells, and the ORF68-3+ cell line failed to produce any infectious virions. Both the ORF68-K435A and 3+ mutant rescue cell lines behaved similarly to wild-type KSHV-infected cells, confirming that defects in virion production are caused by the charge mutations in ORF68, rather than effects on neighboring genes or mutations elsewhere in the viral genome acquired during recombination. Importantly, the K435A and 3+ mutations result in wild-type levels of ORF68 expression (**Fig. 5b**). Interestingly, a small but consistent reduction in the expression level of the late gene K8.1 can be observed in the ORF68.stop, ORF68-K435A, and ORF68-3+ cell lines, suggesting that ORF68 may have other minor roles in gene expression or protein homeostasis during infection (**Supplementary Fig. S7c**).

**Fig. 5.**
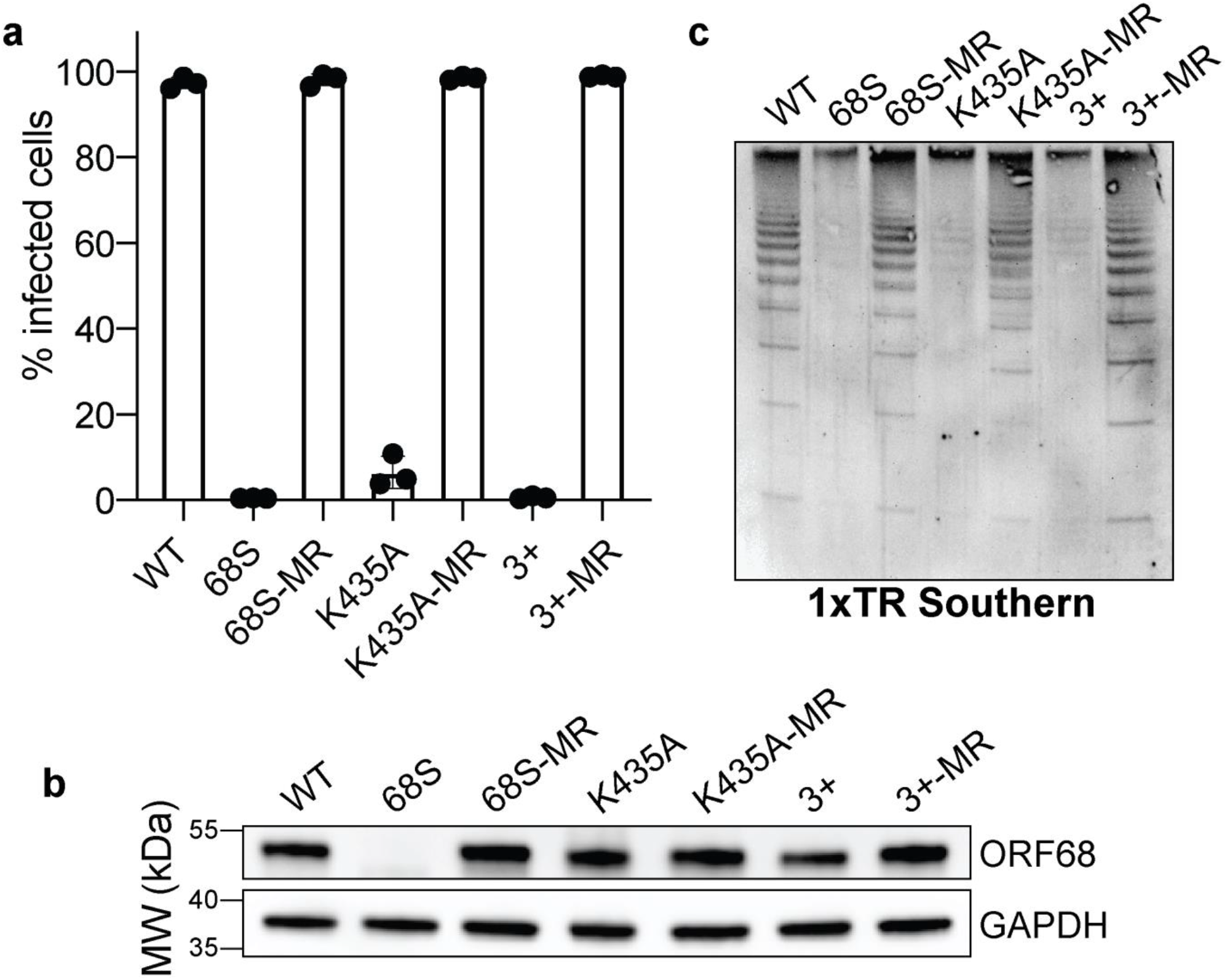
Residues in ORF68 that ablate dsDNA binding *in vitro* are required for genome cleavage and packaging *in vivo*. **a**, iSLK cell lines containing ORF68 mutants (68S, K435A, and 3+) and their corresponding mutant rescues (MR) were established using the KSHV BAC16 system. Progeny virion production by these cell lines was assayed by supernatant transfer and flow cytometry of target cells. **b**, Western blot of whole cell lysate (25 μg) from ORF68.stop iSLK cell lines complemented with pLVX-ORF68 plasmids. GAPDH was used as a loading control. **c**, Southern blot of DNA isolated from iSLK cell lines using a probe for the terminal repeats. DNA was digested with PstI, which cuts within the genome but not within the terminal repeats and generates a ladder of terminal repeat-containing DNA when successful cleavage and packaging occurs.

Given the close association of packaging with replication, we tested if the charge mutations in ORF68 have an effect on DNA replication. We measured viral DNA replication by qPCR and found no difference between the wild-type virus and the ORF68.stop, ORF68-K435A, or ORF68-3+ viruses (**Supplementary Fig. S7d**). While the K435A mutant rescue virus had levels comparable to both wild-type and the K435A mutant, the rescue cell lines for the ORF68 null and 3+ mutants had higher levels of replication, which may be due to changes in the efficiency of cell line establishment.

Using an assay that relies on the intimate coupling between genome packaging and cleavage and that probes the cleavage state of terminal repeat by Southern blot^13^, we previously demonstrated that an ORF68-null virus is defective for viral genome packaging. Given the drastic defects observed in virion production for the K435A and 3+ mutants, while showing no effects on DNA replication, we sought to determine if these mutations act at the step of cleavage and packaging. As we previously demonstrated, the terminal repeats are not cleaved in cells lacking ORF68 (**Fig. 5c**). The ORF68-K435A and 3+ cell lines reveal similarly prominent defects in genome cleavage that are rescued by their respective mutant rescue lines. Thus, the virion production defect observed in the K435A and 3+ cell lines is caused by a failure to properly cleave and package the viral genome, suggesting that the ability of ORF68 to bind nucleic acid through its positively charged central channel is required for successful packaging.

## DISCUSSION

Here, we present the structure of the only functionally undefined component of the herpesviral packaging machinery and reveal that its ability to bind DNA, likely involving its positively charged central channel, is critical for genome packaging. KSHV ORF68 adopts a novel homopentameric ring structure with largely α-helical monomers stabilized by multiple zinc fingers. Comparison to the EBV BFLF1 structure reveals extremely high structural homology, and homology models as well as negative-stain EM analyses of homologs from the further diverged alpha- and beta-herpesviruses indicate a common monomer fold and pentameric quaternary structure. Thus, the core architecture is conserved across herpesviruses from all subfamilies. These structures provide an important framework for mechanistic dissection of the roles that ORF68 and its homologs play in herpesviral packaging.

The importance of ORF68 and its homologs for genome cleavage and packaging has been studied in several herpesviruses, and deletion of these proteins results in a common phenotype, indicative of a conserved function^12–15^. However, our understanding of the packaging process lacks a clear role for this protein^6^ (**Fig. 6a**). UL32, the homolog from HSV-1, has been described as a glycoprotein, yet functional and structural analyses suggest that this is not the case^17,24^. Furthermore, despite earlier work suggesting that UL32 was involved in localizing capsids to replication compartments during packaging, this was not observed upon generation of a virus containing a full deletion of UL32^12,14^. More recent work proposed that UL32 is important for regulation of disulfide bond formation during infection^12^, which was based on the observation that several cysteine-rich motifs are required for infectious virion production, and is consistent with our finding that these cysteines are involved in zinc finger motifs and thus play critical roles for ORF68 stability. Studies in HSV-1 and HCMV demonstrate that the homologs of ORF68 are not involved in expression or localization of the portal or terminase, as deletion of UL32 or UL52, respectively, has no effect on these properties^15,25^. A plethora of mass spectrometry data from different purified herpesviruses, along with recent cryo-EM reconstructions of herpesvirus capsids, suggest that neither ORF68 nor its homologs are packaged into virions. Thus, although it is well established that ORF68 and its homologs are essential for packaging, its role in this process has remained elusive. Given the properties of ORF68 that we have identified – namely, conserved pentameric symmetry with a positively charged channel important for dsDNA binding and virion production – what role could ORF68 be playing during packaging?

**Fig. 6.**
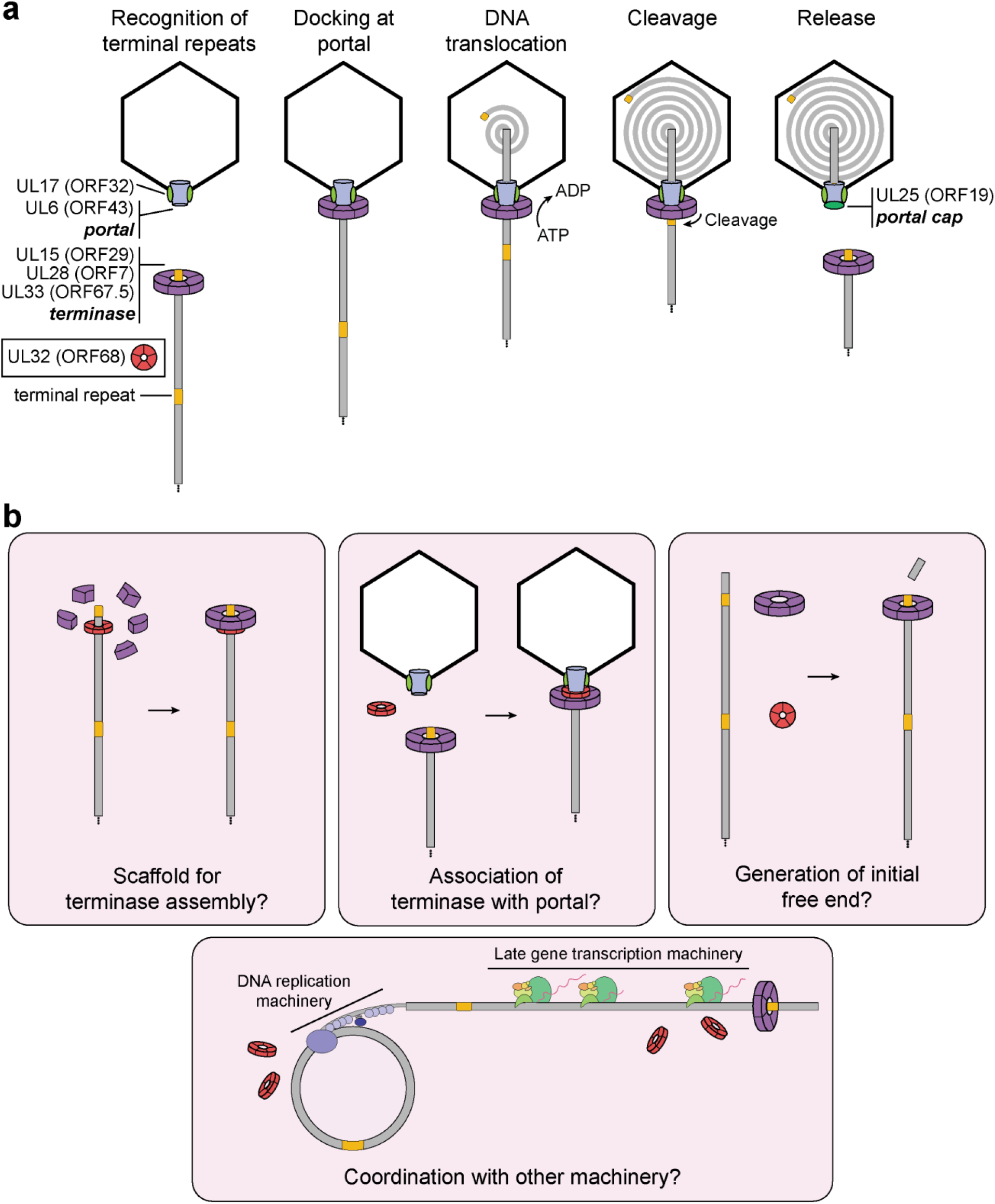
Model of herpesviral cleavage and packaging. **a**, Genes required for cleavage and packaging in HSV-1 and their homologs in KSHV (listed in brackets) are listed. UL15, UL28, and UL33 form the terminase complex that must dock with the portal protein (UL6) and capsid-associated proteins (including UL17). The terminase then translocates the dsDNA genome into the capsid and cleaves within the terminal repeats once a full unit-length genome has been packaged. After release of the remaining genome, the UL25 portal cap binds to stabilize the packaged genome. The role of UL32 (KSHV ORF68) has not been determined. **b**, Possible roles of ORF68 during packaging could include acting as a scaffold for the terminase to bind the nascent genome (left), acting as an adaptor terminase association with the portal (middle), or promoting formation of the initial free end on nascent genomes (right). Further potential roles include interfacing with the DNA replication machinery or late gene transcription machinery.

Similarities between the capsid structure, portal, and terminase in tailed bacteriophages and herpesviruses strongly suggest that these viruses share ancestrally ancient packaging machinery^1^. Although ORF68 has no identifiable structural or functional homolog in phages, several previously studied model phages have accessory factors critical for efficient packaging. These factors highlight possible analogous roles of ORF68 and its homologs in herpesvirus packaging. In phage λ, the factor gpFI is required for efficient packaging^26–28^. gpFI is thought to facilitate binding of the DNA-engaged terminase to the capsid (prohead) through interactions with the major head protein^26,27,29^. Phage phi29 encode a pentameric structural RNA (pRNA) that is sandwiched between the portal and terminase during packaging^30–32^. Although structurally unrelated to gpFI or pRNA, ORF68 by analogy may act as a bridge between the portal and the terminase. Recent reconstructions of the HSV-1 and KSHV capsids^9,11^ revealed their structures at atomic resolution and resolved the portal vertex through which the DNA genome is packaged. The portal cap, which has pentameric symmetry, sits atop this vertex and is thought to be composed of HSV-1 UL25 (KSHV ORF19). Notably, binding of the portal cap to the portal vertex reveals how proteins with pentameric symmetry, like ORF68, could interface with the dodecameric portal.

We propose that ORF68’s role in packaging is to assist docking of the DNA-bound terminase complex with the portal machinery, perhaps by acting as a scaffold that confers five-fold symmetry for the terminase (**Fig. 6b**). Nonspecific DNA binding by ORF68 could promote formation of the terminase-DNA complex and drive the packaging reaction forward. The genome may thread through the channel of the pentameric ring, which in its most constricted region is still wide enough to accommodate dsDNA. Although a recent structure of the terminase complex from HSV-1 revealed hexameric symmetry in solution, its symmetry while bound to the portal remains unknown^33^. It is well-established that phage motors can adopt different stoichiometries, but actively packaging motors are pentameric^32,34,35^. Future work should seek to determine whether ORF68 makes stable or transient protein-protein interactions with components of the packaging machinery and influences assembly or stoichiometry of the terminase complex. Alternatively, ORF68 could be involved in the initial generation of a free dsDNA end for packaging, a process which is poorly understood. It could also be involved in regulation or coordination of packaging with DNA replication and late gene machinery, as coupling between these processes has been observed in phage^36^.

Although ORF68 and its homologs share a conserved structure and likely play similar roles in packaging across the herpesviruses, they are expressed with divergent kinetics, suggesting potential additional functions. HSV-1 UL32 and HCMV UL52 are late genes, consistent with a primary role in packaging^12,15^, whereas KSHV ORF68 is an early gene, accumulating prior to the expression of several capsid proteins^13^. It remains to be determined whether ORF68 has additional roles early in infection. We demonstrated that deletion of ORF68 can be partially complemented by other gammaherpesvirus homologs (mu68 and BFLF1). Interestingly, deletion of the ORF68 homolog in the alphaherpesvirus pseudorabies virus (PrV) can be partially complemented by expression of HSV-1 UL32; however, the inverse complementation was not observed, indicating that some functions or interactions are sufficiently similar for PrV virion production, but not HSV-1 virion production^37^. PrV UL32 and HSV-1 UL32 share ~52% sequence similarity, as do ORF68 and EBV BFLF1 or MHV mu68, suggesting that in both cases common residues or structural motifs mediate shared interactions to carry out critical functions in the cell. Future high-resolution structural studies of other homologs may identify shared and divergent features, and it will be interesting to investigate how homologs lacking the third zinc finger motif (i.e. HCMV UL52) are structurally stabilized.

Despite widespread prevalence of herpesviruses and the significant diseases they cause, there is no cure for any herpesvirus, and a vaccine exists only for the alphaherpesvirus varicella zoster virus (VZV). The majority of antiherpesviral drugs target the DNA replication machinery, but have several disadvantages, including the emergence of resistance mutations and a narrow spectrum of use^38–40^. However, Letermovir, a recently FDA-approved drug for the treatment of HCMV in stem cell transplant recipients, targets the HCMV terminase through an unknown mechanism^41–43^. Letermovir lacks activity against other herpesviruses^44^, suggesting that key differences exist in their conserved machinery. Our work represents a major advance in understanding the complexities of herpesviral DNA packaging and the mechanistic role of one of its essential components. Additional work will be required to elucidate the detailed role of ORF68 and its homologs in packaging. Understanding the molecular underpinnings of packaging and the differences in mechanism across the alpha, beta, and gammaherpesviruses is critical for the future development of antiherpesvirals as effective therapeutics.

## MATERIALS AND METHODS

### Plasmids

ORF68 was amplified from pcDNA4/TO-ORF68-2xStrep13 and mu68 was amplified from MHV68 BAC DNA45. UL52 was amplified from HCMV Towne DNA. UL32 was amplified using nested PCR from HSV-1 KOS DNA. ORF68 and its homologs were subcloned into the NotI and XhoI sites of pcDNA4/TO-2xStrep (N-terminal) using InFusion cloning (Clontech) (Addgene #x-x). Mutations in ORF68 (Addgene #x-x) and UL32 (Addgene #x-x) were generated using inverse PCR site-directed mutagenesis with Phusion DNA polymerase (New England Biolabs) with primers as listed in **Supplementary Table 3**. PCR products were DpnI treated, ligated using T4 PNK and T4 DNA ligase and transformed into *Escherichia coli* XL-1 Blue cells.

The expression plasmid for ORF68 was previously described (Addgene #x) and is a pHEK293 UltraExpression I vector (pUE1-TSP) (Clontech) which encodes an N-terminal Twin-Strep tag and the coding region for ORF68 including its native start codon, separated by an HRV 3C protease cleavage site. This plasmid was used as a template for inverse PCR to generate linearized pUE1-TSP vector containing the Twin-Strep tag and HRV 3C site. BFLF1, UL32, and mutants of ORF68 were subcloned from their respective pcDNA4/TO vectors into linearized pUE1-TSP vector using InFusion cloning to generate expression constructs (Addgene #x-x).

Plasmids for lentiviral transduction (pLJM1-2xStrep or untagged wild-type ORF68, mutants, and homologs) (Addgene #x-x) were generated by subcloning into the AgeI and EcoRI sites of pLJM1 modified to confer resistance to zeocin (Addgene #19319) using InFusion cloning. Lentiviral packaging plasmids psPAX2 (Addgene plasmid #12260) and pMD2.G (Addgene plasmid #12259) were gifts from Didier Trono.

### Transfections

HEK293T cells were plated and transfected after 24 h at 70% confluency with PolyJet (SignaGen) or polyethylenimine (PEI). Cells were harvested after 24 h (for expression studies in **Fig. 2**) or 48 h (for large-scale protein expression). For analysis of protein expression, cells were washed with PBS, pelleted at 1,000 x g for 5 min at 4°C, and lysed by resuspension in lysis buffer (150 mM NaCl, 50 mM Tris-HCl pH 7.4, 1 mM EDTA, 0.5% NP-40, and protease inhibitor [Roche]) with rotation at 4°C for 30 min. Lysates were clarified by centrifugation at 21,000 x g for 10 min at 4°C. Lysate (33 μg) was used for SDS-PAGE and Western blotting in Tris-buffered saline and 0.2% Tween 20 (TBST) using Strep-Tag II HRP (1:2,500; EMD Millipore) and rabbit anti-vinculin (1:1,000; Abcam). Following incubation with primary antibodies, membranes were washed with TBST and imaged (for Strep-Tag II HRP) or incubated with goat anti-rabbit-HRP (1:5,000; Southern Biotech).

### Protein expression and purification

Purification of Twin-Strep tagged ORF68 and BFLF1 for use in crystallography and cryo-EM was performed as described previously^13^. Briefly, pUE1-TSP-ORF68 or BFLF1 was transfected into ~70% confluent HEK293T cells using PEI. Cells were harvested after 48 h and frozen at −80°C. Cells were lysed in lysis buffer (300 mM NaCl, 100 mM Tris-HCl pH 8.0, 5% glycerol, 1 mM DTT, 0.1% CHAPS, 1 μg/mL avidin, cOmplete, EDTA-free protease inhibitors [Roche]), rotated at 4°C for 30 min, sonicated to reduce viscosity, then centrifuged at 50,000 x g for 30 minutes at 4°C. The lysate was filtered through a 0.45 μm filter then purified using Strep-Tactin XT resin (IBA) in running buffer (300 mM NaCl, 100 mM Tris-HCl pH 8.0, 5% glycerol, 0.1% CHAPS, 1 mM DTT). Protein was eluted in running buffer containing 50 mM biotin, concentrated using a 30 kDa cutoff spin concentrator (Millipore), then the 2xStrep tag was removed by cleavage with HRV 3C protease (Millipore) overnight. Protein was further purified by size exclusion over a HiLoad 16/600 Superdex 200 pg column (GE Healthcare) in 100 mM NaCl, 50 mM HEPES pH 7.6, 5% glycerol, 1 mM TCEP-HCl.

The native start methionine was included in the construct, and interestingly we reproducibly saw a ~50% distribution between translation initiation at the methionine before the Twin-Strep tag and at the native methionine for both ORF68 and BFLF1, but not UL32. Little untagged ORF68 or BFLF1 is lost during purification as one Twin-Strep tag in the pentamer is sufficient for enrichment on StrepTactin resin. Proteins used for electrophoretic mobility shift assays and UL32 used in negative stain EM were purified as above, except that 0.5% CHAPS was used during lysis, 1 mM TCEP-HCl was used in lieu of DTT throughout the purification, the 2xStrep tag was not removed by incubation with HRV 3C protease (Millipore), and proteins were not purified by size exclusion chromatography.

### Negative stain and cryo-electron microscopy grid preparation, data collection, image processing, initial model building, and structure determination

For preparation of negative-stain EM grids, ORF68, BFLF1, and UL32 were diluted to ~100-200 nM in Dilution Buffer (60 mM HEPES pH 7.6, 100 mM NaCl, 10 mM MgCl_2_, 0.5 mM TCEP) and stained with 2% uranyl formate (pH 5.5– 6.0) on thin carbon-layered 400 mesh copper grids (EMS)46. Micrographs were collected on a Tecnai 12 microscope (ThermoFisher) operated at 120 keV with 2.2 Å per pixel using a 4k TemCam-F416 camera (TVIPS): 131, 79, and 100 total micrographs for ORF68, BFLF1, and UL32 datasets, respectively. Micrographs were CTF corrected using CTFFIND4^47^ in Relion^48^. Single particles were automatically selected using Gautomatch^49^: 57,510, 29,021, and 29,098 total particles for ORF68, BFLF1, and UL32 datasets, respectively, and 2D classification was performed in Relion^48^.

Cryo-EM grids were prepared by applying 3.5 μL of 5 μM (pentamer) ORF68 (50 mM HEPES pH 7.6, 50 mM NaCl, 50 mM KCl, 5% glycerol, and 1 mM TCEP) and 3.5 μL of 14 μM BFLF1 (60 mM HEPES pH 7.6, 100 mM NaCl, 50 mM KCl, 10 mM MgCl^2^, 0.5 mM TCEP, 0.05% NP-40) to glow-discharged C-Flat holey carbon grids (CF-2/1-3C-T, EMS). The samples were plunge-frozen using a Vitrobot (ThermoFisher) and imaged on a Talos Arctica TEM operated at 200 keV (ThermoFisher). Dose-fractionated imaging was performed by automated collection methods using SerialEM^50^. Data were collected as described in **Supplementary Table 1**. Whole-frame drift correction was performed via Motioncor2^51^ with dose weighting applied.

ORF68 processing: Micrographs were CTF corrected using GCTF^52^ in Relion^48^. 662,435 single particles were automatically selected using Gautomatch^49^, from which 274,167 particles were selected from 2D class averages, generated in Relion. The 3D classification scheme is detailed in **Supplementary Fig. 1**. The final model was refined with C5 symmetry (pentamer) and sharpened using postprocessing to an estimated 3.37 Å. An alanine backbone was modeled manually in *Coot* ^53^, to be used for phasing the crystal structure.

BFLF1 Processing: Micrographs were CTF corrected using CTFFIND4^47^ in Relion^48^. A small subset of micrographs was used to generate initial 2D averages using the Relion Laplacian autopicker, which were then used as a template on the total dataset for template-based particle picking, resulting in 278,234 total particles. After a round of 2D classification, 155,574 particles were selected. The 3D classification scheme is detailed in **Supplementary Fig. 1**. Both CtfRefinement and Bayesian polishing were applied to the final 38,908 particle set, and then refined with D5 symmetry (a decamer of stacked pentamers) and sharpened using postprocessing^48^. The final structure was refined to an estimated 3.60 Å.

A homology model of BFLF1 was generated using the ORF68 atomic model in SWISS MODEL^54^. Subsequent refinement was done with iterative rounds of manual model building in *Coot*^53^ and automated refinement in phenix.refine^55–57^. Data collection and refinement statistics are listed in **Supplementary Table 1**.

### Crystallization and structure determination

Crystals of ORF68 were obtained by hanging drop vapor diffusion with 2 μL of concentrated protein (20 mg/mL) and 2 μL of crystallization solution (310 mM CaCl^2^, 95 mM HEPES pH 7.5, 26.6% PEG-400, 5% glycerol) with equilibration against 1 mL of crystallization solution at 20°C. Crystals grew over the course of 1-3 days and were harvested and flash-cooled in liquid nitrogen without additional cryoprotection. X-ray diffraction data were collected at 100 K on a single crystal at beamline 8.3.1 at the Advanced Light Source (Berkeley, CA). Data were integrated with XDS^58^ and the space group was determined using *POINTLESS*^59^. Data were merged and elliptically truncated to correct for anisotropy using the STARANISO server^60^. The diffraction limits along the a*, b*, and c* axes were 2.86, 2.60, and 2.21 Å, respectively. The corrected anisotropy amplitudes were used for molecular replacement in PHASER^61^ using the partially built cryo-EM model of ORF68. Subsequent refinement was done with iterative rounds of manual model building in *Coot*^53^ and automated refinement with TLS in phenix.refine^55–57^. Data collection and refinement statistics are listed in **Supplementary Table 2**.

All figures were generated with PyMol (http://www.pymol.org). The electrostatic surface was calculated using APBS^62^ in PyMol. Surface conservation as depicted in **Supplementary Fig. S3** was generated using the ConSurf server^63^. Homology models for UL32 and UL52 were generated using SWISS-MODEL^54^.

### Sequence alignment

The sequence of ORF68 was used for a BLAST search to find homologs in the *Herpesviridae*. No clear homologs were readily identified in the *Herpesvirales* families *Alloherpesviridae* and *Malacoherpesviridae*. A multiple sequence alignment was generated using Clustal Omega64 and manually edited to condense long insertions relative to ORF68.

### Cell lines

HEK293T cells were maintained in DMEM supplemented with 10% FBS (Seradigm). HEK293T cells constitutively expressing ORF68 (HEK293T-ORF68) were previously described^13^ and were maintained in DMEM supplemented with 10% FBS and 500 μg/ml zeocin. iSLK-puro cells were maintained in DMEM supplemented with 10% FBS and 1 μg/ml puromycin. The iSLK-BAC16 system^23^ consists of the KSHV genome on a bacterial artificial chromosome BAC16 and a doxycycline-inducible copy of the KSHV lytic transactivator RTA. All iSLK-BAC16 cell lines were maintained in DMEM supplemented with 10% FBS, 1 mg/mL hygromycin, and 1 μg/ml puromycin. iSLK-BAC16-ORF68.stop cells were complemented with pLJM1-2xStrep- or untagged ORF68 wild-type, mutants, or homologs by lentiviral transduction as described below and were maintained in DMEM supplemented with 10% FBS, 1 mg/mL hygromycin, 1 μg/ml puromycin, and 500 μg/ml zeocin.

### Cell line establishment and viral mutagenesis

Complemented iSLK-BAC16-ORF68.stop cells^13^ were generated by lentiviral transduction. Lentivirus was generated in HEK293T cells by co-transfection of pLJM1-ORF68 wild-type, mutant, or a homolog along with the packaging plasmids pMD2.G and psPAX2. After 48 h, the supernatant was harvested and syringe-filtered through a 0.45 μm filter. The supernatant was diluted 1:2 with DMEM and polybrene was added to a final concentration of 8 μg/ml. 1 x 106 freshly trypsinized iSLK-BAC16-ORF68.stop cells were spinfected in a 6-well plate for 2 h at 876 x g. After 24 h the cells were expanded to a 10 cm tissue culture plate and selected for 2 weeks in DMEM supplemented with 10% FBS, 1 mg/mL hygromycin, 1 μg/ml puromycin, and 500 μg/ml zeocin.

All viral ORF68 mutants were generated using the scarless Red recombination system in BAC16 GS1783 *Escherichia coli* as previously described^23^. Modified BACs were purified using a Nucleobond BAC 100 kit (Clontech). BAC quality was assessed by digestion with RsrII and SbfI (New England Biolabs). Latently infected iSLK cell lines with modified virus were generated by transfection of HEK293T-ORF68 cells with 5 μg BAC DNA using PolyJet reagent (SignaGen). The following day, transfected HEK293T cells were trypsinized and mixed 1:1 with freshly trypsinized iSLK-puro cells and treated with 30 nM 12-O-tetradecanoylphorbyl-13-acetate (TPA) and 300 μM sodium butyrate for 4 days to induce lytic replication. iSLK cells were then selected in medium containing 300 μg/ml hygromycin B, 1 μg/ml puromycin, and 250 μg/ml G418. The hygromycin B concentration was increased to 500 μg/ml and 1 mg/ml until all HEK293T cells died.

### Virus characterization

For reactivation studies, 1 x 10^6^ iSLK cells or iSLK.ORF68.stop cells complemented with wild-type or mutant ORF68 were plated in 10 cm dishes for 16 h, then reactivated with 1 μg/ml doxycycline and 1 mM sodium butyrate for an additional 72 h or left unreactivated. Infectious virion production was determined by supernatant transfer assay. Supernatant from reactivated iSLK cells was syringe-filtered through a 0.45 μm filter, then 2 mL was spinoculated onto 1 x 10^6^ freshly trypsinized HEK293T cells for 2 h at 876 x g. After 24 h, the media was aspirated, the cells were washed with cold PBS and crosslinked in 4% PFA (Electron Microscopy Services) diluted in PBS. Cells were pelleted at 1,000 x g for 5 minutes at 4°C, resuspended in PBS, and 50,000 cells/sample were analyzed on a BD Accuri 6 flow cytometer. The data were analyzed using FlowJo version 10.

To determine fold DNA replication, reactivated and unreactivated iSLK cells were rinsed with PBS, scraped, pelleted at 1,000 x g for 5 min at 4°C, and stored at −80°C. Cells were resuspended in 600 μL of PBS, of which 200 μL was purified using a NucleoSpin Blood kit (Macherey Nagel) according to the manufacturer’s instructions. The fold DNA replication was quantified by qPCR using iTaq Universal SYBR Green Supermix (Bio-Rad) on a QuantStudio3 Real-Time PCR machine. DNA levels were quantified with primers specific for the KSHV ORF57 promoter and normalized to human CTGF promoter and to unreactivated samples to determine fold replication (**Supplementary Table 3**).

Total protein was isolated from reactivated iSLK cells at 72 h. Samples were resuspended in lysis buffer, rotated for 30 min at 4°C, and clarified by centrifugation at 21,000 x g for 10 min at 4°C. Lysate (25 μg) was used for SDS-PAGE and Western blotting in Tris-buffered saline and 0.2% Tween 20 (TBST) using rabbit anti-K8.1 (1:10,000), rabbit anti-ORF68 (1:5,000), rabbit anti-ORF6 (1:10,000), and mouse anti-GAPDH (1:1,000; Abcam). Rabbit anti-K8.1, anti-ORF6, and anti-ORF68 were previously described^13,65^. Following incubation with primary antibodies, membranes were washed with TBST and incubated with goat anti-rabbit-HRP (1:5,000; Southern Biotech) or goat anti-mouse-HRP (1:5,000; Southern Biotech).

### Southern blotting

iSLK-BAC16 cells (1 x 10^6^) were harvested after 72 h of reactivation. Cells were rinsed with PBS, scraped, pelleted at 1,000 x g for 5 min at 4°C, and stored at −80°C. Cells were resuspended in 700 μL of Hirt lysis buffer (10 mM Tris-HCl pH 7.4, 10 mM EDTA, 0.6% SDS) and incubated at room temperature for 5 min. NaCl was added to a final concentration of 0.85 M and samples were rotated at 4°C overnight. The following day, insoluble material was pelleted by centrifugation at 21,000 x g for 30 min at 4°C. The supernatant was then treated with 100 μg/mL RNase A (ThermoFisher) for 1 h at 55°C then with 200 μg/mL proteinase K (Promega) for 1 h at 55°C. DNA was isolated by phenol-chloroform extraction and ethanol precipitation. DNA (5 μg) was digested with PstI-HF (New England Biolabs) overnight then separated by electrophoresis on a 0.7% agarose 1x TBE gel. The gel was denatured in 0.5 M NaOH, 1.5 M NaCl, neutralized in 1 M Tris pH 7.4, 1.5 M NaCl, then transferred to an Amersham Hybond-N+ membrane (GE Healthcare) by capillary action in 20x SSC (3M NaCl, 0.3 M sodium citrate pH 7.0) overnight, and cross-linked to the membrane in a StrataLinker 2400 (Stratagene) using the AutoUV setting. The membrane was treated according to the DIG High Prime DNA Labeling and Detection Starter Kit II (Roche) according to the manufacturer’s instructions using a DIG-labeled DNA probe corresponding to a single repeat of the KSHV terminal repeats as previously described^13^. After overnight hybridization, washing, blocking, incubation with anti-DIG-AP antibody, the membrane was visualized on a ChemiDoc MP imaging system (Bio-Rad).

### Electrophoretic mobility shift assays

Fluorescein-labeled dsDNA probes (**Supplementary Table 3**) (Integrated DNA Technologies) were prepared as 2x stocks (20 nM) in binding buffer (100 mM NaCl, 20 mM Hepes pH 7.5, 5% glycerol, 0.05% CHAPS). Wild-type or mutant ORF68 (retaining the 2xStrep tag and HRV 3C protease cleavage site) was diluted in binding buffer containing 0.2 mg/mL BSA. Final binding reactions were prepared by mixing equal volumes of probe DNA and protein, and thus contained 10 nM DNA probe, 100 mM NaCl, 20 mM Hepes pH 7.5, 5% glycerol, 0.05% CHAPS, 0.1 mg/mL BSA, and variable concentrations of ORF68. Concentrations listed are for the monomer. Samples were incubated at room temperature for 20 minutes prior to electrophoresis on a 5% polyacrylamide (29:1 acrylamide:bis-acrylamide)/1x Tris borate gel at 2W at 4°C. Gels were imaged using a ChemiDoc MP imaging system (Bio-rad). Results were analyzed using Fiji^66^. The percent bound probe was determined by dividing the intensity of the shifted band by the total intensity of the lane. Binding curves were generated using non-linear regression in GraphPad Prism 8 to the Hill equation: % bound = (Bmax *[protein]^H)/(*K*d^H + [protein]^H).

### Data availability

Atomic coordinates and structure factors for ORF68 have been deposited in the Protein Data Bank with accession code 6XF9. Diffraction images have been deposited in the SBGrid Data Bank under ID 794 (https://doi:10.15785/SBGRID/794). Cryo-EM maps for ORF68 and BFLF1 have been deposited in the Electron Microscopy Data Bank with accession codes EMD-22167 and EMD-22168. The atomic model of BFLF1 was deposited in the Protein Data Bank with accession code 6XFA. Final coordinate sets, structure factors with calculated phases, and cryo-EM maps are provided as **Supplementary Data 1**.

## ACKNOWLEDGEMENTS

We thank current and former members of the Glaunsinger and Martin labs, particularly Angelica Castañeda, for their helpful suggestions and critical reading of the manuscript. We are grateful to Eric Montemayor for advice. We thank beamline staff at ALS 8.2.2 and 8.3.1 for assistance with data collection, and UC Berkeley Cal-Cryo Facility members for assistance with cryo-EM data collection. A.D. is the Rhee Family Fellow of the Damon Runyon Cancer Research Foundation (DRG-2349-18). S.G. is a Howard Hughes Medical Institute Fellow of the Damon Runyon Cancer Research Foundation (DRG-2342-18). B.G. and A.M. are investigators of the Howard Hughes Medical Institute. This research was supported by NIH R01AI122528 to B.G. Beamlines 8.2.2 and 8.3.1 of the Advanced Light Source, a U.S. DOE Office of Science User Facility under Contract No. DE-AC02-05CH11231, are supported in part by the ALS-ENABLE program funded by the National Institutes of Health, National Institute of General Medical Sciences, grant P30 GM124169-01.

## AUTHOR CONTRIBUTIONS

A.D., M.G., and L.S. purified protein. A.D. and S.G. built atomic models. A.D. generated and characterized cell lines and performed *in vitro* binding assays. B.G. and A.M. supervised research. All authors edited and approved the paper.

## SUPPLEMENTARY FIGURES

**Supplementary Figure S1.**
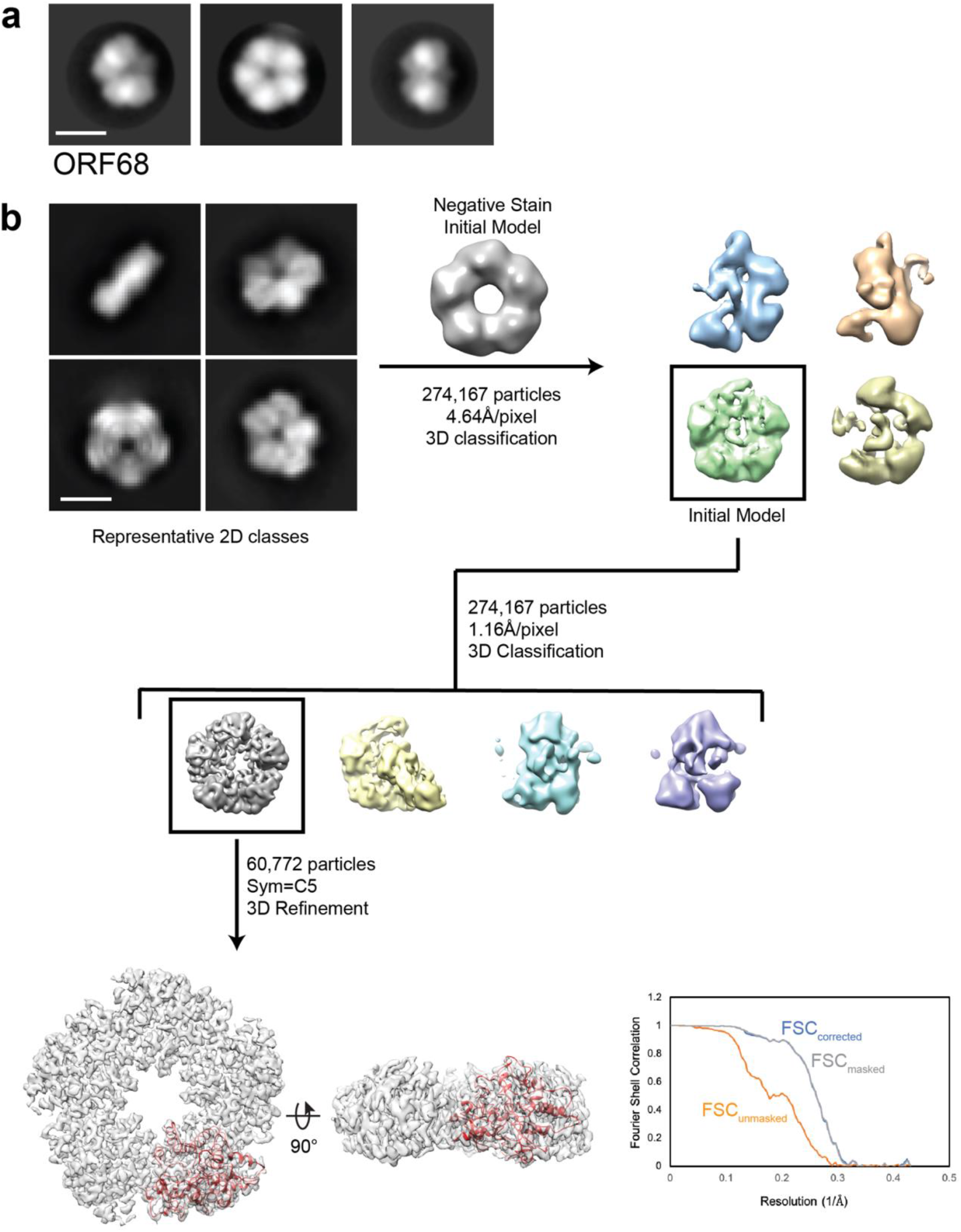
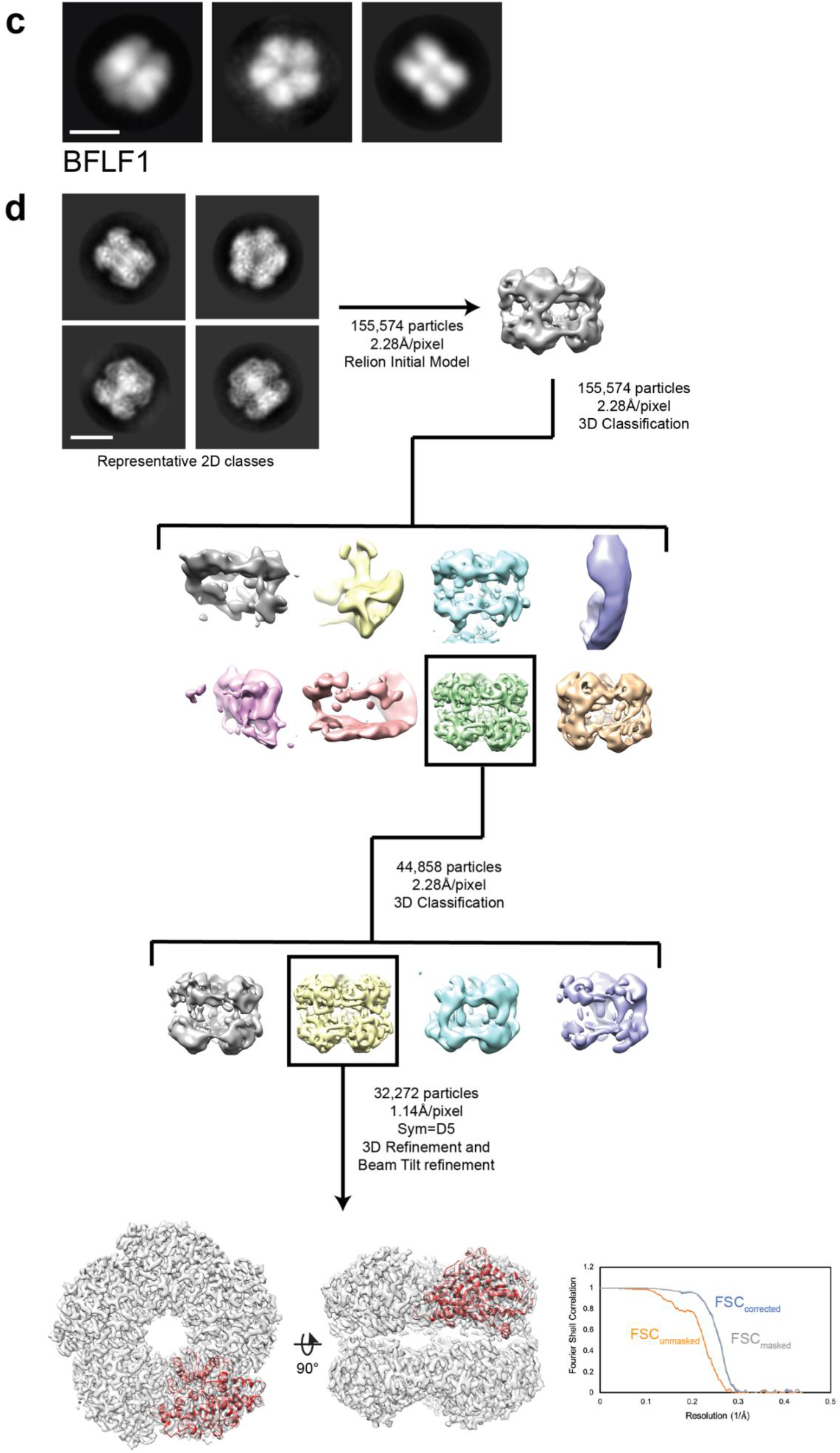
EM data analysis. **a**, Representative negative stain 2D class averages of ORF68. Scale bar =100 Å. **b**, Representative 2D class averages and data-processing workflow for single-particle cryo-EM analyses of ORF68. The best class was refined to an estimated 3.37 Å resolution, as indicated by the depicted FSC plot. Scale bar = 100 Å. **c**, Representative negative stain 2D class averages of BFLF1. Scale bar =100 Å. **d**, Representative 2D class averages and data-processing workflow for singleparticle cryo-EM analyses of BFLF1. The best class was refined to an estimated 3.60 Å resolution, as indicated by the FSC plot. Scale bar = 100 Å.

**Supplementary Figure S2.**
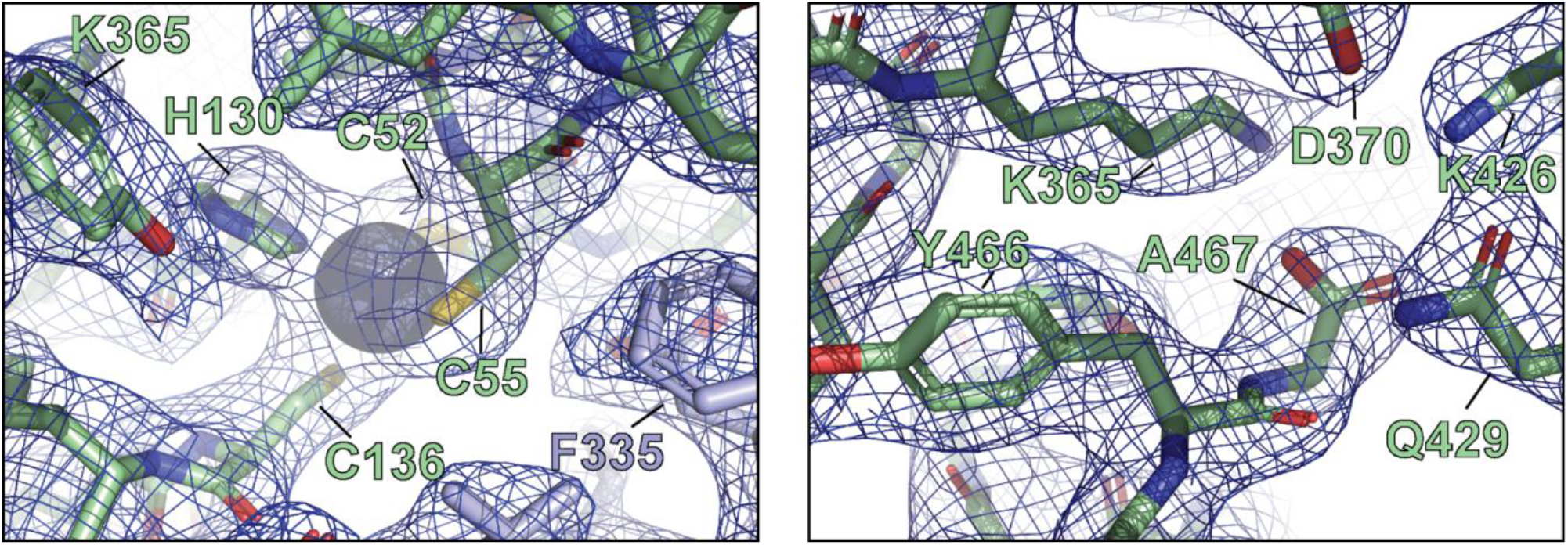
Representative electron densities and X-ray structural details of ORF68. Representative local 2*m*F_o_-*D*F_c_ maps are contoured at 1σ.

**Supplementary Figure S3.**
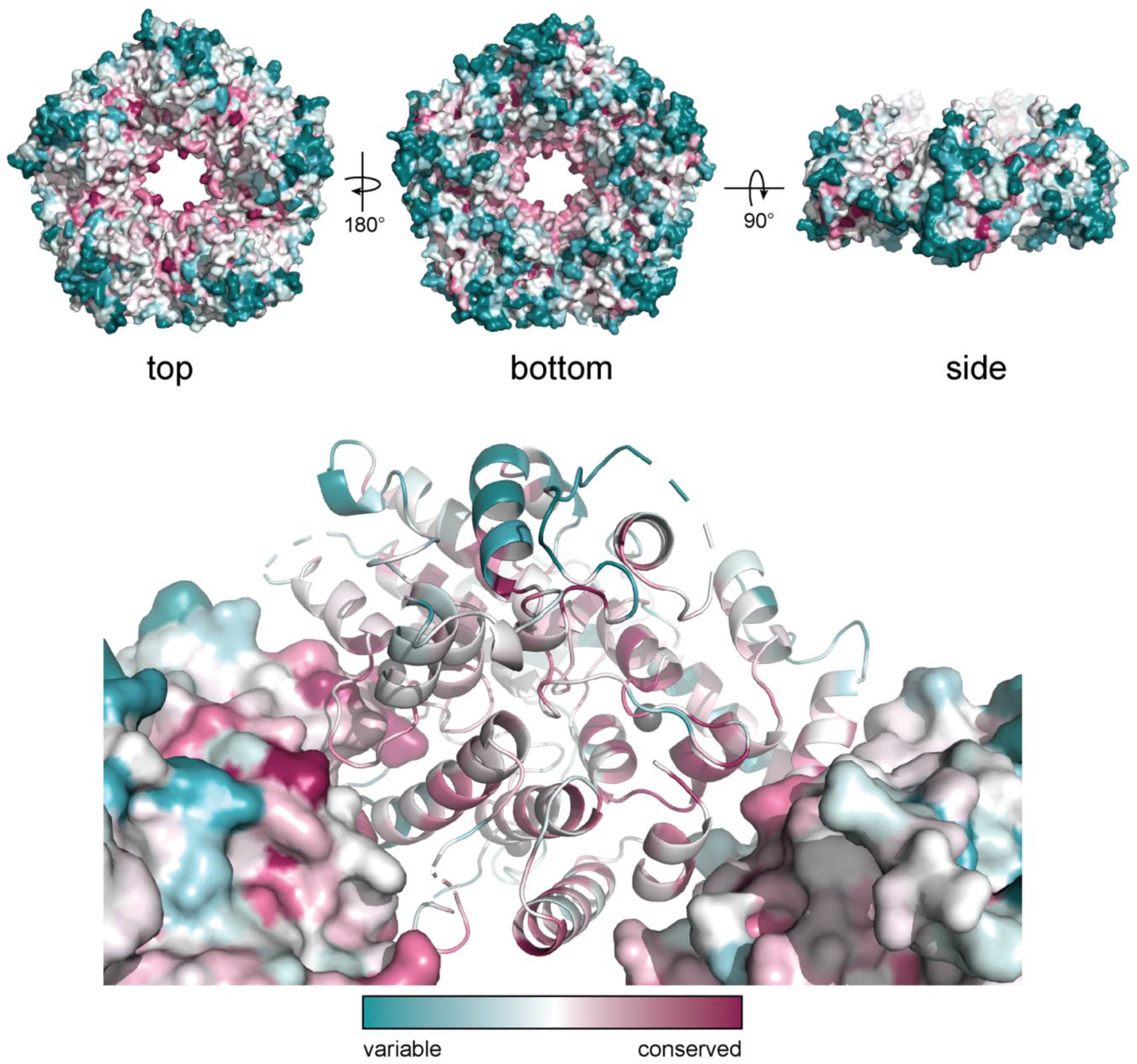
ConSurf model of ORF68. Conservation of residues within ORF68 as calculated by the ConSurf server63, where blue residues are variable (nonconserved) and red are conserved.

**Supplementary Figure S4.**
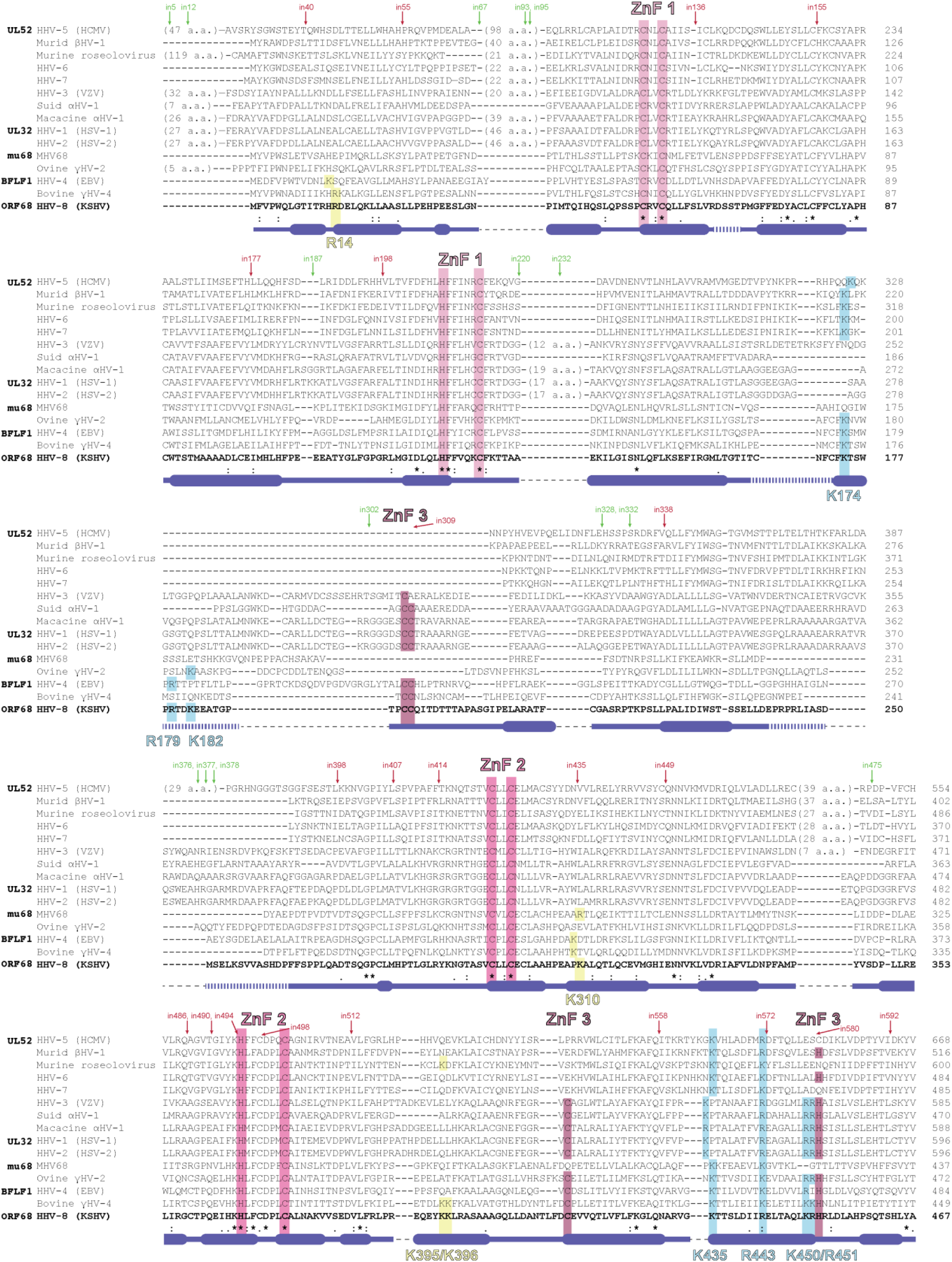
Sequence alignment of homologs of ORF68. Homologs of ORF68 were identified using BLAST and aligned using Clustal Omega, then manually edited to condense long insertions relative to ORF68. (*) indicates perfect conservation, (:) indicates strong conservation, and (.) indicates weak conservation. Secondary structure as observed in the ORF68 structure is indicated under the alignment, with thick lines for α-helices, thin lines for unstructured regions or loops, and dotted lines for disordered regions of the model. Thin dotted black lines indicate an insertion in the alignment relative to ORF68. The residues involved in the three zinc finger (ZnF) motifs are highlighted in shades of pink. Positively charged residues selected for mutation located in the pore of the ring are highlighted in blue, while residues selected for mutation outside of the pore are highlighted in yellow. A transposon screen of UL32 (the ORF68 homolog from HSV-1) was previously performed67 and revealed insertion sites (“in#”) within the protein that were either tolerant (labeled in green) or intolerant (labeled in red) of insertion of a 5-amino acid transposon, as assessed by complementation of a UL32-null virus.

**Supplementary Figure S5.**
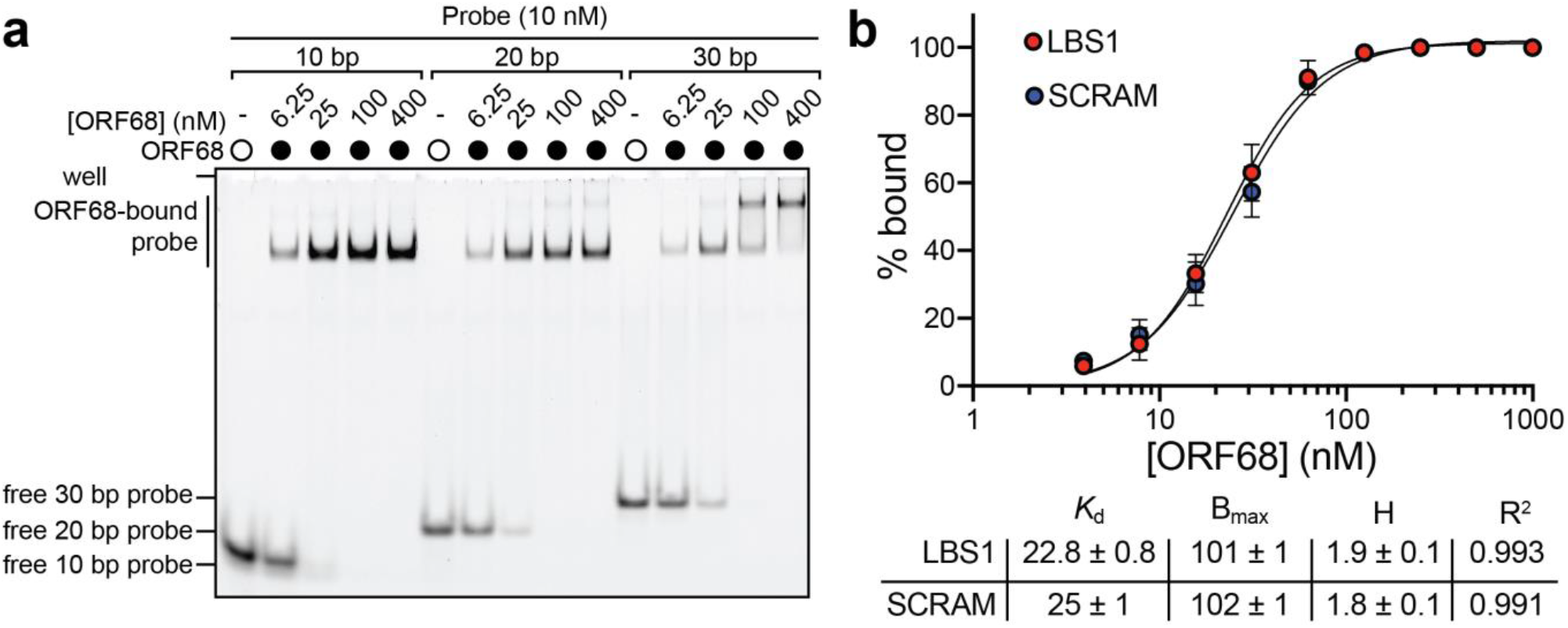
ORF68 non-specifically binds nucleic acid. **a**, Native gel for the electrophoretic mobility shift assay of ORF68 binding to 5’-fluorescein labeled dsDNA probe with 10, 20, or 30 bp. **b**, Binding curves for a 85% GC-rich probe (LBS1) and a 50% GC-rich probe (SCRAM) (top). Data represent the mean ± s.d. of three independent experiments. Data were fit with a nonlinear regression to the Hill equation, with best fit derived binding parameters within the 95% CI (bottom): *K*_d_ (binding affinity), B_max_ (maximum specific binding), H (Hill coefficient), and R_2_ (goodness of fit).

**Supplementary Figure S6.**
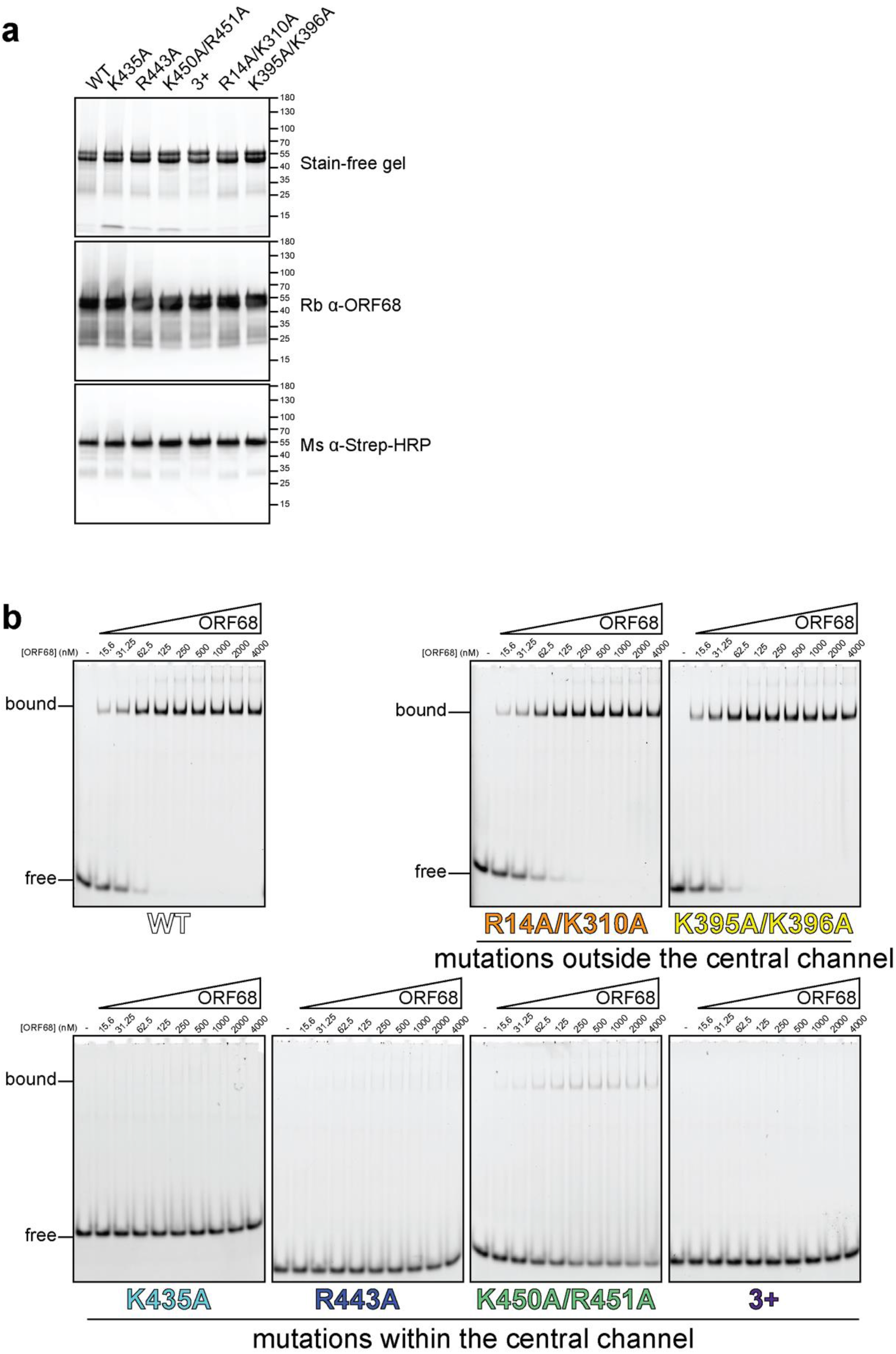
ORF68 mutants can be purified, but mutations in the central channel prevent dsDNA binding. **a**, SDS-PAGE gel of purified recombinant wild-type and mutant ORF68 variants (7.5 μg) used for dsDNA-binding assays. Proteins were visualized by stain-free imaging (top), followed by western blotting for ORF68 (middle), and the Strep tag (bottom). **b**, Representative native gels for electrophoretic mobility shift assays with wild-type or mutant ORF68 and a 20 bp dsDNA probe.

**Supplementary Figure S7.**
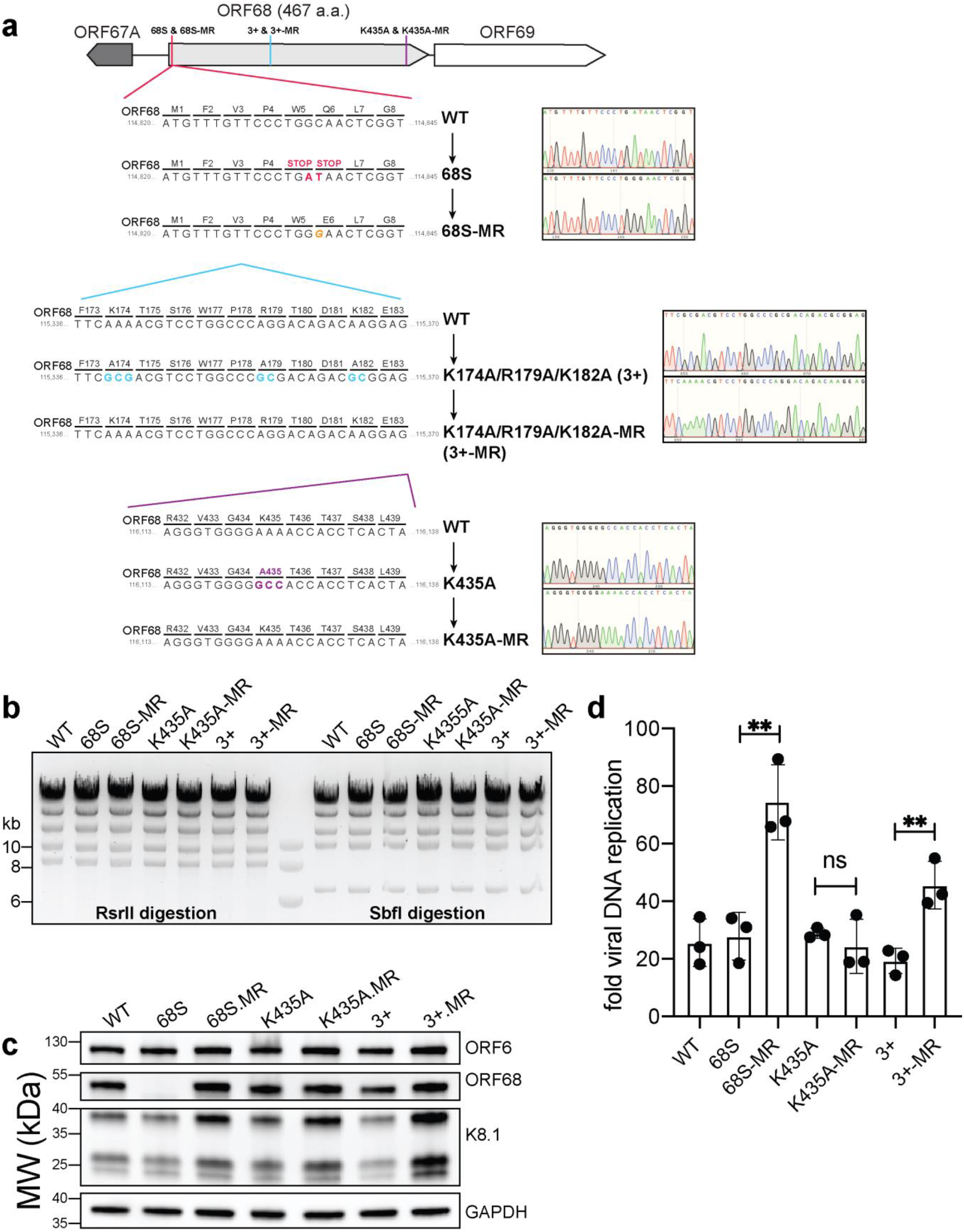
Construction and validation of mutant viruses. **a**, Schematic of the genomic locus of ORF68, with the location of introduced mutations depicted in detail below. Sanger sequencing traces for the mutants and corresponding mutant rescues are shown to the right. **b**, Digestion of recombinant BACs with RsrII and SbfI was used to assess whether large-scale recombination had occurred during mutagenesis. **c**, Western blot of whole cell lysate (25 μg) from ORF68.stop iSLK cell lines. GAPDH was used as a loading control. ORF6 is an early gene and K8.1 is a late gene. **d**, Viral DNA replication was measured by qPCR before and after reactivation. Data are from three independent biological replicates, with statistics being calculated using an unpaired *t* test. **, p<0.01.

**Supplementary Table S1.** Cryo-EM data collection statistics.

|  | **ORF68** | **BFLF1** |
| --- | --- | --- |
| Microscope | Talos Arctica | Talos Arctica |
| Nominal Mag. |  |  |
| Detector | K2 | K3 |
| Pixel Size (Å/pixel) | 1.16 | 1.14 |
| Exposure (s) | 8 | 2.4 |
| Frame rate (s) | 0.509 | 0.05 |
| Total electron dose | 48 | 48 |
| Defocus Range (µm) | -1.0 to -2.5 | -1.5 to -3.0 |
| Total Micrographs | 2408 | 839 |
| Total Particles | 662,435 | 278,234 |

The cryo-EM maps for ORF68 and BFLF1 and the coordinate set for BFLF1 are available in **Supplementary Data File 1**.

**Supplementary Table S2.** X-ray data collection and refinement statistics for ORF68.

| Parameter | ORF68 (PDB XXXX) |
| --- | --- |
| **Data collection statistics** |  |
| Wavelength (Å) | 1.11589 |
| Resolution range (Å) | 49.54 - 2.22 (2.48 - 2.22) |
| Space group | C222<sub>1</sub> |
| Unit cell dimensions |  |
| a, b, c (Å) | 135.0 224.0 192.3 |
| $\alpha=\beta=\gamma$ (°) | 90 |
| Total reflections | 1,213,852 (51,624) |
| Unique reflections | 91,387 (4,570) |
| Multiplicity | 13.3 (11.3) |
| Completeness (%) |  |
| spherical | 63.7 (11.3) |
| ellipsoidal | 95.3 (79.0) |
| Mean $I/\sigma(I)$ | 19.4 (1.7) |
| Wilson B-factor | 54.8 |
| R-merge | 0.100 (1.715) |
| R-meas | 0.104 (1.796) |
| R-pim | 0.028 (0.525) |
| CC<sub>1/2</sub> | 0.999 (0.565) |
| **Refinement statistics** |  |
| R<sub>work</sub>/R<sub>free</sub> | 0.22/0.25 (0.38/0.35) |
| Number of non-hydrogen atoms | 16,391 |
| macromolecules | 16,373 |
| ligands | 15 |
| solvent | 3 |
| Protein residues | 2124 |
| RMS (bonds) | 0.013 |
| RMS (angles) | 1.38 |
| Ramachandran favored (%) | 98.07 |
| Ramachandran allowed (%) | 1.93 |
| Ramachandran outliers (%) | 0.00 |
| Rotamer outliers (%) | 0.33 |
| Clashscore | 11.40 |
| Average B-factor | 62.79 |
| macromolecules | 62.81 |
| ligands | 48.61 |
| solvent | 40.49 |
| Number of TLS groups | 22 |

Statistics for the highest-resolution shell are shown in parentheses. The STARANISO server was used for ellipsoidal truncation<sup>60</sup>. The worst diffraction limit after cut-off was 2.99 Å. The ellipsoidally truncated data set was deposited in the Protein Data Bank and is available in **Supplementary Data File 1**. Merged diffraction data that has not been ellipsoidally truncated is also available in **Supplementary Data File 1**. The coordinate set deposited in the Protein Data Bank is available as **Supplementary Data File 1**.

**Supplementary Table S3.**
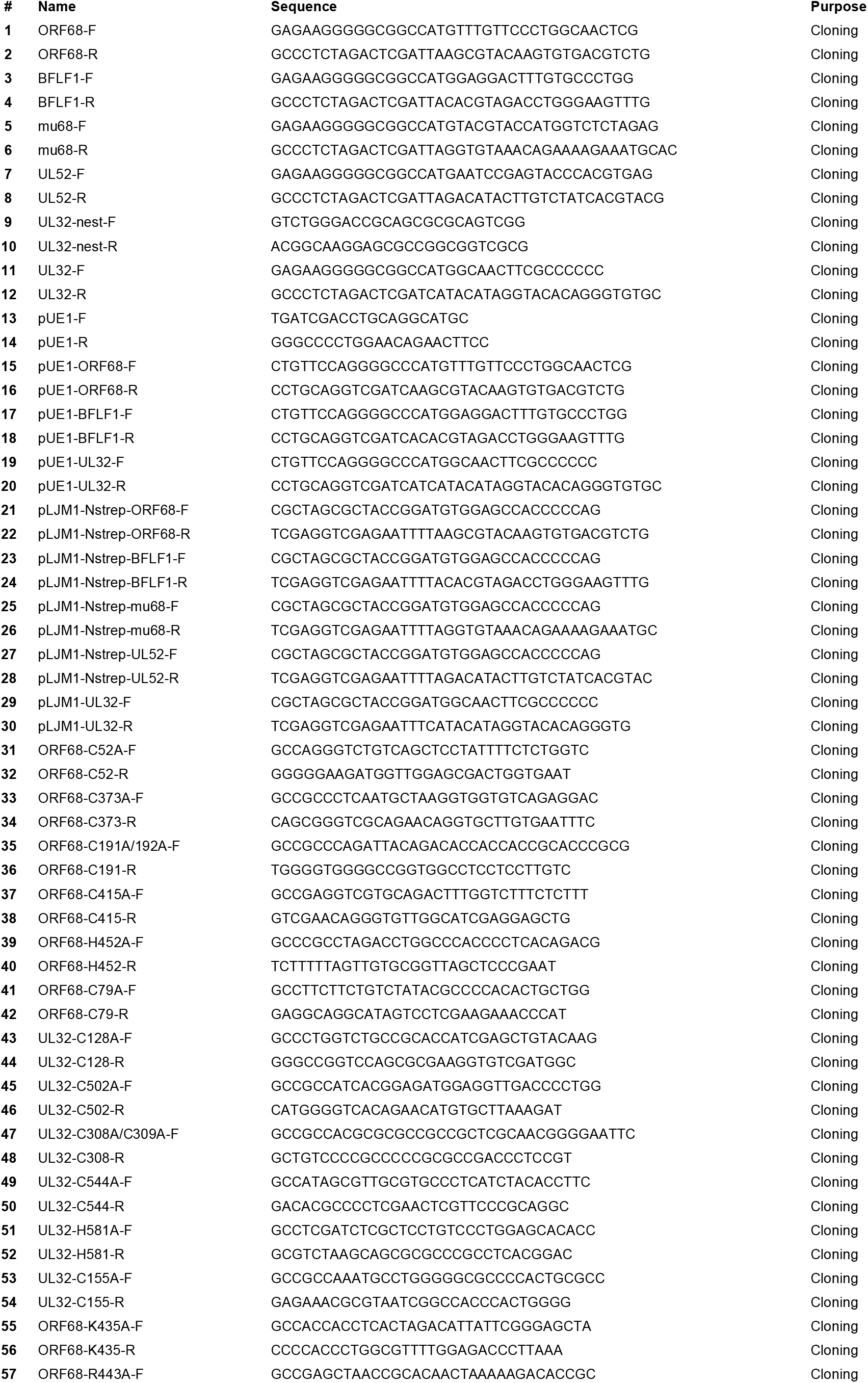
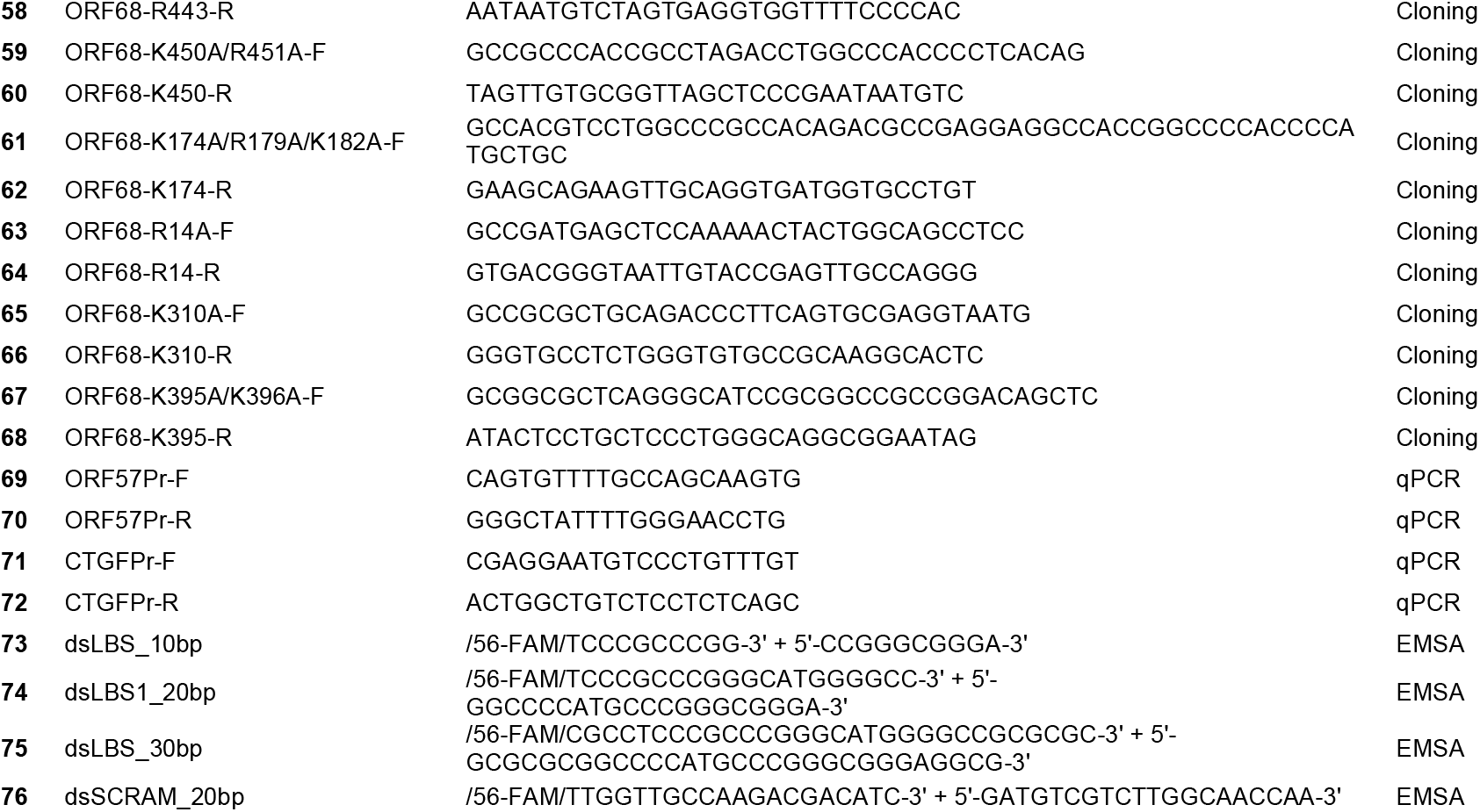
Oligonucleotides used for cloning, qPCR, and EMSAs.

